# Modeling transcriptional regulation of the cell cycle using a novel cybernetic-inspired approach

**DOI:** 10.1101/2023.03.21.533676

**Authors:** Rubesh Raja, Sana Khanum, Lina Aboulmouna, Mano R. Maurya, Shakti Gupta, Shankar Subramaniam, Doraiswami Ramkrishna

## Abstract

Quantitative understanding of cellular processes, such as cell cycle and differentiation, is impeded by various forms of complexity ranging from myriad molecular players and their multilevel regulatory interactions, cellular evolution with multiple intermediate stages, lack of elucidation of cause-effect relationships among the many system players, and the computational complexity associated with the profusion of variables and parameters. In this paper, we present an elegant modeling framework based on the cybernetic concept that biological regulation is inspired by objectives embedding entirely novel strategies for dimension reduction, process stage specification through the system dynamics, and innovative causal association of regulatory events with the ability to predict the evolution of the dynamical system. The elementary step of the modeling strategy involves stage-specific objective functions that are computationally-determined from experiments, augmented with dynamical network computations involving end point objective functions, mutual information, change point detection, and maximal clique centrality. We demonstrate the power of the method through application to the mammalian cell cycle, which involves thousands of biomolecules engaged in signaling, transcription, and regulation. Starting with a fine-grained transcriptional description obtained from RNA sequencing measurements, we develop an initial model, which is then dynamically modeled using the cybernetic-inspired method (CIM), utilizing the strategies described above. The CIM is able to distill the most significant interactions from a multitude of possibilities. In addition to capturing the complexity of regulatory processes in a mechanistically causal and stage-specific manner, we identify the functional network modules, including novel cell cycle stages. Our model is able to predict future cell cycles consistent with experimental measurements. We posit that this state-of-the-art framework has the promise to extend to the dynamics of other biological processes, with a potential to provide novel mechanistic insights.

**STATEMENT OF SIGNIFICANCE:** Cellular processes like cell cycle are overly complex, involving multiple players interacting at multiple levels, and explicit modeling of such systems is challenging. The availability of longitudinal RNA measurements provides an opportunity to “reverse-engineer” for novel regulatory models. We develop a novel framework, inspired using goal-oriented cybernetic model, to implicitly model transcriptional regulation by constraining the system using inferred temporal goals. A preliminary causal network based on information-theory is used as a starting point, and our framework is used to distill the network to temporally-based networks containing essential molecular players. The strength of this approach is its ability to dynamically model the RNA temporal measurements. The approach developed paves the way for inferring regulatory processes in many complex cellular processes.

## INTRODUCTION

Cellular processes involve a complex network of molecular interactions associated with response to external stimuli, signal transduction, chromatin modifications, transcriptional regulation and a host of regulatory mechanisms. Painstaking biochemical analyses on these complex regulatory processes have provided some insights although incomplete. Further, there is little data on the kinetics of regulatory processes even at a coarse-grained level making quantitative modeling of the cellular processes difficult (1–3). The ability to infer all the regulatory players and processes is out of scope of currently available experiment methods and it is essential to decipher an approach that will be able to account for the regulatory processes in an implicit manner. Longitudinal time-series data from high throughput measurements provide details about potential regulation but incorporating such vast data into mathematical models has proven to be a challenge. Previously established modeling approaches, including steady-state stoichiometric models (4), kinetic models (1, 5, 6), dynamic-FBA models (7), and discrete Boolean models (8), have faced these challenges. Constraint-based approaches such as FBA (4) and network decomposition approaches like Elementary Modes (9) can aid in modeling regulatory mechanisms. Most of these represent static or steady-state situations and some with pseudo-steady state perspectives that are very useful for reactions with disparate rates. However, the dynamic complexity of eukaryotic cell behavior makes pseudo-steady state assumptions void.

Cybernetic method (10), which inherits its name from optimizing models with goals, where regulatory processes are inferred implicitly through parameters, has been successful in dynamically modeling bacterial metabolism whose objective is cell growth optimality (10–13). Recently, we extended this approach to modeling macrophage lipid metabolism with the goal of maximal expression and activation of inflammatory cytokine TNFα (14, 15). With this single objective function, we were able to incorporate regulatory mechanisms involved in prostaglandin metabolism. In all these examples only one stage with a single well-defined objective function is assumed making it difficult to extend it to multi-stage processes with a distinct objective function associated with each stage.

The mammalian cell cycle is an exemplary cellular process (16) where cells replicate through a complex set of events across multiple stages and it illustrates the complexity associated with such biological processes, namely stage-specific regulation, and stage-specific phenotype endpoints, i.e., biological objectives for each stage. It is important to understand that the ‘stages’ mentioned here represent periods of time within which the cell is expected to have a distinct objective. While ‘stages’ are not the same as cell cycle ‘phases’, they can coincide with each other. The cell cycle goes through different experimentally defined phases (G1, S, G2, and M), where during the G1 phase the cell increases its cellular contents and grows, followed by the S phase where the chromosomes are duplicated, with growth and preparation for the mitosis in the G2 phase followed by mitosis (cell division) in the M phase prior to returning to the G1 phase (3, 17, 18). Extant dynamical modeling approaches are not capable of addressing multiple stages with distinct and multiple objective functions (18–20).

We report in this manuscript a novel cybernetic-inspired method (CIM) framework that accounts for stage-specific modeling with distinct objectives where each stage involves implicit modeling of regulatory processes involved at that stage. In the cell cycle modeling using a set of fine-grained time-series transcriptomic measurements (3), we develop stage-specific models with specific objectives, and model their dynamical behavior controlled by explicitly unspecified regulatory processes. It is appropriate that we model the transcriptomic data as the stages in the cell cycle are transcriptionally-driven. The biological objective of such a system can be stated as the optimization of cell cycle where the objective is different at each stage depending on the specific functional differences across each stage. During each stage, the objective is mathematically represented to maximize the weighted production rates of all the transcripts where the weights are expected to implicitly depend on the functional importance of the genes during that stage. For each stage representing a different objective, we modeled them by having different weights where these weights are inferred by fitting to the data.

For most biological systems like the cell cycle, there is limited availability of information relating to the numerous interactions among the myriad components and the amount of data needed to estimate the model parameters (21, 22). The longitudinal RNA-seq data in the form of time-series is a great resource for developing and validating our CIM framework for dynamical modeling. The temporal molecular data have significant correlations both at the same time point and at distinct time points reflecting cause-effect relationships. To infer this notion of correlation and crosstalk between different RNA transcripts, we explored the power of information-theoretic approaches (23) by relating the components that have a higher degree of mutual information. Mutual information, an information-theoretic approach, detects non-linear correlations between datasets and can be used to formulate the causal interactions (24, 25). Moreover, causality is fundamental to the cybernetic approach in order that the system tweaks the relevant variables to realize objectives. Specifically, we used the time-delayed mutual information (TDMI) method to identify the causal relations and temporal correlations between any two RNA transcripts within the network, and by setting a threshold we selected a preliminary network to model with our CIM framework (Figure 1). Mutual information thus plays a very key enabling role in the success of the model framework.

**Figure 1:**
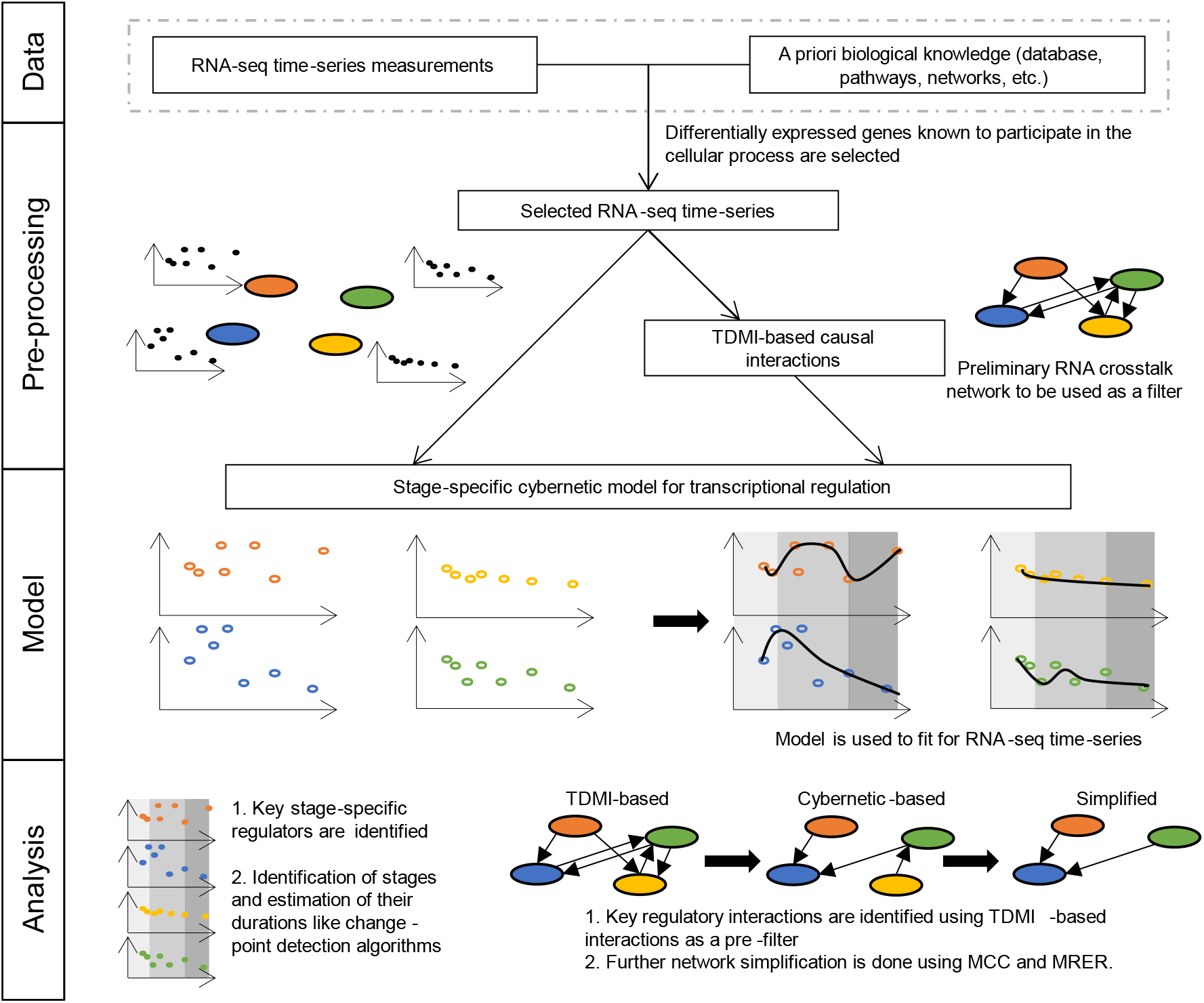
Schematic for cybernetic-inspired method (CIM). Using the selected subset of RNA-seq measurements, a preliminary RNA crosstalk network is developed using the time-delayed mutual information (TDMI). The stage-specific cybernetic model is then applied on the data and the TDMI-network. By optimizing the parameters, the data is fitted, and the key species and regulatory interactions are identified. This approach also identifies the stage durations. Further network simplification is carried out using the Maximal Clique Centrality (MCC) and the Maximal RNA expression rate (MRER). (All data and networks shown in this figure are for representation and not real.)

Further, the stages with distinct objective functions during cell cycle are not pre-defined. While it is natural to match the stages with experimentally derived temporal regimes like cell cycle phases, each of these phases may subsume other undefined stages. To correctly identify stages, we used the CIM to simultaneously work as change-point detector while fitting the RNA-seq data. This approach provided key descriptions of regulatory interactions, especially in identifying the key stage-specific genes. Since matching observations of system variables with their model counterparts is the manner in which the weights (and other model parameters) are determined, they represent the system choice arising from the built-in biological objectives.

## METHODS

### Identification of causal interactions using time-delayed mutual information

The advantages of working with information-theoretic approaches (26) are that they facilitate quantifying the interactions between datasets and avoid assuming a functional form of their relationship (27). Specifically, we employed mutual information (MI), an information-theoretic approach that quantifies the linear and non-linear interactions between the variables (28–31).

The mutual information *I*(*X*; *Y*) for random variables *X* and *Y*, given random samples {*y*_1_,…, *y*_S_} and {*x*_1_,…, *x*_S_} (S denotes the number of samples), with joint probability function *p*(*x_i_*, *y_j_*) and marginal probability functions *p_x_*(*x_i_*) and *p_y_*(*y_j_*) is:

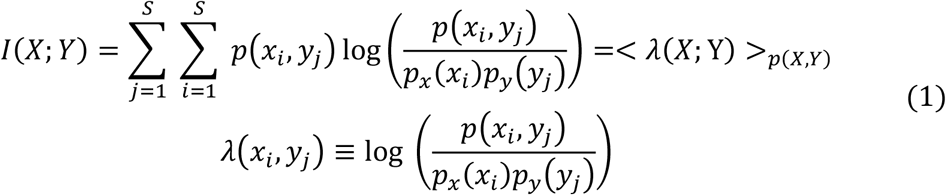

where < . > denotes the average operator. *I*(*X*; *Y*) is the average of the difference between the log likelihood of two variables log *p*(*x_i_*, *y_j_*) and single variable log *p_x_*(*x_i_*), log *p_y_*(*y_j_*) with respect to joint probability density *p*(*x_i_*, *y_j_*).

In our approach, the series *X_t_* and *Y_t_* represent RNA time-series data. The goal was to calculate mutual information *I*(*X_t_*; *Y_t_*) for different combinations of RNA time-series measurements, e.g., *I*(*RNA*_1_, *RNA*_2_). The above MI definition assumes that the samples *x_i_* are sampled independently, i.e., {*x*_1_,…, *x*_n_ belongs to an independent set (the same is true for *y_j_* samples as well). However, for time-series data, the assumption of independence will be invalid due to temporal correlations between data points at various times.

Galka et al. developed the “innovation approach to mutual information” for temporally correlated time series (32). While the authors derived an *I*(*X_t_*; *Y_t_*) formula for a Gaussian distributed innovation set that forms an independent and identically distributed (iid) sequence, here, we developed an approach valid for all distributions. For time series, *λ*(*x_i_*, *y_j_*) in equation (1) was redefined by pairing the time points as:

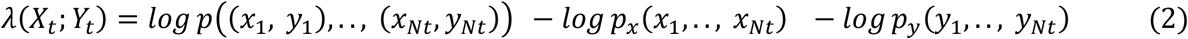

Equation (2) had high-dimensional joint distributions with complicated structure. To simplify the structure of these distributions we described these correlations by the corresponding optimal predictors of *X_t_* and *Y_t_* using time-series prediction model: auto-regression (AR) or vector auto-regression (VAR). The ‘lag’ hyperparameter for the AR and VAR models were selected based on minimization of AIC. The calculated residuals (*ϵ_t_*) based on the expected model (*E*) are called “innovations”.

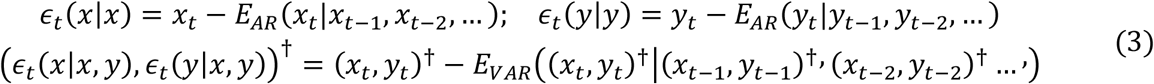

Since, the probabilities of original series are same as the probabilities of their innovations (32) equation (2) becomes:

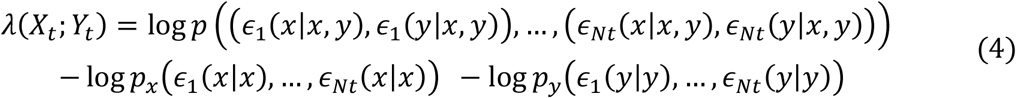

For optimal predictors of time-series, the innovations are white noise, and they are independent. Moreover, if the time points are assumed to form an independent and identically distributed sequence, the joint probability of the innovation time series deciphers as the product of marginal probability densities. The equation (4) became:

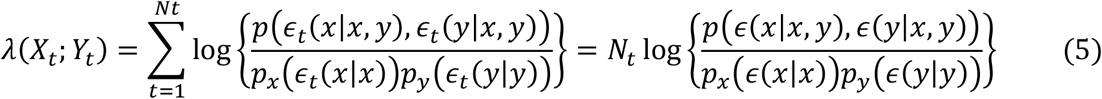

The mutual information per time point was then be calculated using equations (1) and (5) as follows:

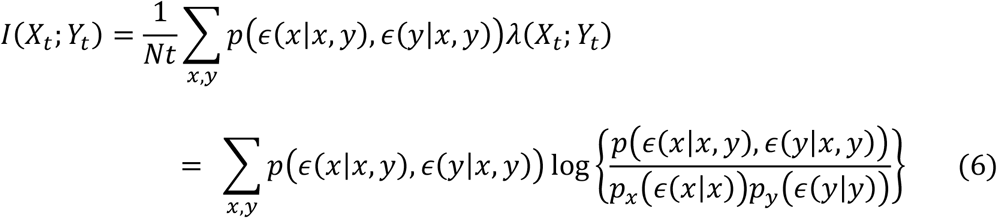

The probabilities of the above equation was estimated using kernel density estimators (KDE) after finding the innovations using auto-regression and vector auto-regression models (33). We then normalized the mutual information value by dividing it with average of the entropies: *I_norm_*(*X_t_*; *Y_t_*) = 2*I*/(*X_t_*; *Y_t_*/(*H*(*x_t_*) + *H*(*y_t_*)).

To identify the causal interactions between any two species within the network, we used the time-delayed mutual information (TDMI). For calculating TDMI, a relative time-delay (τ) was introduced between the series *X_t_* and *Y_t_*, and then the mutual information formula is applied to this *τ*-shifted time series, i.e., *X_t_* and *Y_t–τ_*. Thus, the TDMI calculation entails evaluating mutual information as a function of *τ*. Further, at the maximum MI value (*TDMI^max^*), the sign of *τ* can be used to infer the direction of causality between *X_t_* and *Y_t_* processes from the information transfer perspective (34, 35).

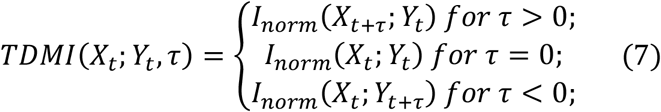

### Cybernetic-inspired method (CIM) for transcription regulation

Cybernetic models in the past have been predominantly used in modeling metabolic regulation where two sets of control variables for enzyme activation and synthesis control the dynamics based on a system objective. Here, we specifically modeled RNA expression levels where the underlying regulation is implicitly incorporated in the cybernetic model using the new control variables ‘*u*’ and ‘*v*’ that are different from the past and will be defined below. The transcriptional regulation can be affected by the expression of RNAs, histones, transcription factors, or chromatin modifications and topological constraints associated with the state. In order to incorporate the effect of crosstalk or interactions across RNA players due to these multi-level intermediates, we defined a lumped species ‘*g*’ called the regulator of gene expression. Thus, in our formulation, the RNA expression depends on the species ‘*g*’ and ‘*g*’ depends on the indirect interactions (proxied through other RNAs). Here are the model equations:

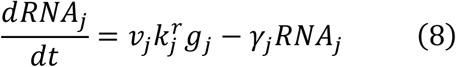

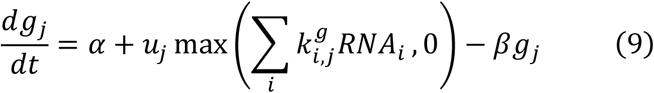

In equation (8), the parameter 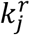 is the *RNA_j_* expression rate constant and *γ_j_* is the *RNA_j_* degradation rate. In equation (9), the three parameters denote the basal priming rate *α*, the interaction parameter for *RNA_j_* by *RNA_i_* 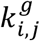, and the decrease of priming level by the rate constant *β*. The interaction parameter was set to be non-zero only for possible interactions pre-determined using a priori knowledge or data-driven approach. They are either activating (+) or repressing (-) rate constant allowing the term 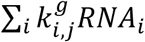 in equation (9) to become negative for some conditions. We allowed a basal transcription rate (*g_j_* = *α*/*β*) even when the above term becomes negative. We implemented this constraint by using max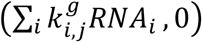 in equation (9). The control variables ‘*u*’ and ‘*v*’ are regulating the level and strength of priming for the RNA transcription respectively and are defined based on the cybernetic objective that is either intuitively described or phenotypically described from the experimental data.

We defined that the objective of this system is to maximize 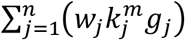, that is, to maximize the sum of weighted production rates of the RNAs. This objective form is similar to those used in the original cybernetic models and means that the RNA production is optimized based on the functional requirement decided by the weights. The control variables, *v_j_* and *u_j_*, are computed by solving the optimal control problem resulting in the Proportional and Matching laws (10), respectively, as follows:

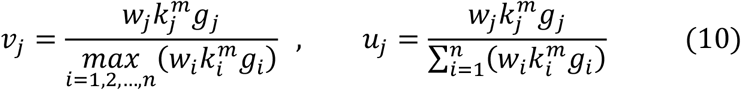

The expressions for the control variables, though appearing simplistic, make the ODE system (equations (1) and (2)) to be non-linear.

In this formulation, we attached time/age to the cell transition. If the values *t*_1_, *t*_2_, *t*_3_, *t*_4_, *t*_5_ and *t*_6_ represent the times at which the cell’s objective changes for *G*1 → *S*_1_, *S*_1_ → *S*_2_, *S*_2_ → *S*_3_, *S*_3_ → *G*2, *G*2 → *M* transitions and cell division, respectively, then the weights are as follows:

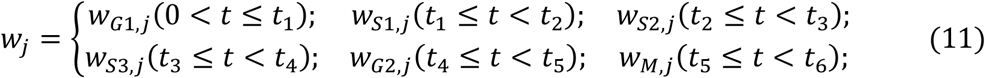

The fits are attained based on the global minimization of the sum of the squares of normalized fit errors (SSE) where the RNA-specific fit errors are normalized by dividing it with their maximum value.

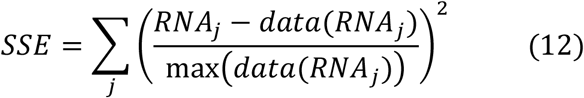

### Application of CIM framework to mouse cell cycle measurements

Transcriptomic time-series measurements for synchronized cell cycle of mouse embryonic fibroblast were available (17) from our prior work. The mouse embryonic fibroblast (MEF) cells were serum-starved to arrest the cell cycle and the serum was added at time 0 to simultaneously initiate and synchronize the cell cycle (17). RNA-seq data was measured at 96 different time points covering more than one full cell cycle (3, 17). Of the 4248 genes differentially expressed (more than 2-fold up or down as compared to t=0) at one or more time points, we selected 63 canonical cell cycle genes and 23 more transcription factor genes (a total of 86) based on our prior model (3).

## RESULTS

### Modeling transcriptomic regulation during the cell cycle

We illustrate the versatility of our CIM framework by applying it to model transcriptional regulation during cell cycle progression in a mammalian cell. While our CIM approach has ability to infer the regulatory interactions on unbiased choice of molecules, it is important to start with a model-network based on a priori knowledge or by time-series analysis techniques to avoid model overparameterization (Figure 1) (36, 37). This will also enable the approach to relate to known biology. Using the initial set of 86 RNA transcripts as nodes (see Methods for selection criteria), we evolved the network by introducing mechanistic causality using our longitudinal measurements and mutual information.

### Causal interaction network development using time-delayed mutual information

We refined our preliminary network model using the time-delayed mutual information (TDMI; see Methods). We applied our approach to the time series data of the 86 selected genes to calculate the TDMI for the range of delays (*τ*) between −20 to 20 hours. In Figure 2, we show a sample calculation for TDMI between the two nodes (genes) in the network, Tgfb1 and Ets1 (Figures 2A and 2B). We used the “innovation approach” (32) for TDMI calculation in which we calculated the residuals using AR and VAR models (see Methods). Then, we estimated the probability functions based on mono-variate and bi-variate KDE models (Figure 2C, D, and E) (33). We then estimated the TDMI across different *τ* values (see Methods; Figure 2F). We can see that the TDMI peaks at a positive lag value of 4 hours (Figure 2F) implying that Ets1 is the cause and Tgfb1 the effect. We then repeated the calculation for every pair of the selected genes. We selected the top 0.1 percentile of the interactions by comparing the max value of TDMI for each pair (Figures 2G and S1). These interactions among the RNA transcripts (nodes) are visualized as networks using Cytoscape (Figure 2G). Though these selected interactions are highly possible, this TDMI-based approach does not provide any mechanistic proof for these interactions. So, we need a mechanistic model like the CIM approach to further validate the network.

**Figure 2:**
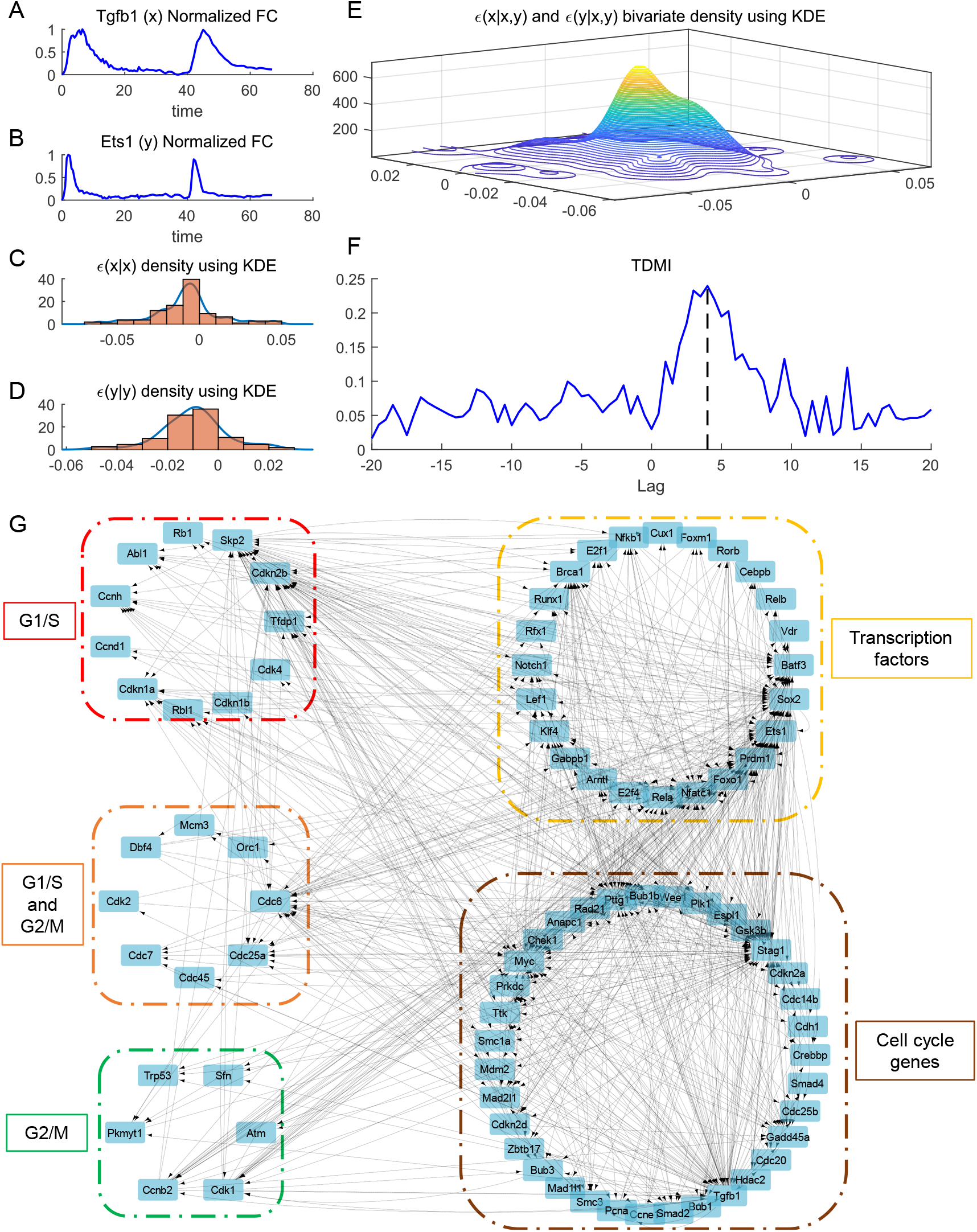
Estimating time-delayed mutual information (TDMI) using the “innovation” approach and using it to generate a causal interaction network. (A,B) Time-series RNA transcriptomics measurements for Tgfb1 and Ets1. (C,D) Mono-variate KDE for probability densities of residuals (E) Bi-variate KDE for probability density of residuals. (F) TDMI calculated between time-series of Tgfb1 and Ets1 for different lags. (G) Interaction network development for the cell cycle model using TDMI. Network construction is based on TDMI threshold of top 0.1 percentile. The boxed sections are color-coded based on category mentioned.

### Stage-specific CIM for transcriptional regulation

We developed CIM framework to model transcriptional regulation incorporating stage-varying and multifactorial regulation (see Methods). During cell cycle, the objective or goal of the system depends on the current phenotype of the cell and needs to be redefined each time the cell undergoes a major transition (stage-change) owing to transcriptional remodeling. While a single multi-weighted objective function describes regulation for short periods within a single stage, for modeling long time-series measurements, we had to incorporate stage-specific objective functions. We mathematically defined the objective for each stage is to maximize the sum of weighted production rates of the RNAs where the weights are computationally fitted (see Methods). The importance of an RNA during a particular stage can be inferred based on its weight for the stage. The higher the weight for RNA, the higher the functional necessity for it during a particular stage and vice versa, and thus this approach connects the mathematical objective to biological objective. During cell cycle, we incorporated the changes in the objective functions by fitting for different weights for each stage. Each stage was modeled with its own *W_j_* represented by *w*_k,*j*_, where k = {*G*1, *S*_1_, *S*_2_, *S*_3_, *G*2, *M*} represents the sequential objective-changing stages of the cell. It is important to note that while most cell cycle phases (experimentally-defined) can be modeled using a specific single objective (e.g., *G*1, *G*2, *M*), some stages will require special modifications based on the biological process. Here, during S phase, there is competition between transcription events and DNA replication, and there is a global anti-correlation between replication and transcription timing making it impossible to model using a single objective (38). This observation is an important part of the model development here which necessitates a stage-specific objective approach and thus a significant departure from cybernetic modeling of microbial metabolism. Therefore, S phase can have multiple stages and here we used 3 stages to model S phase (represented as *S*_1_, *S*_2_, *S*_3_), with each stage having its own objective (Figure 3).

**Figure 3:**
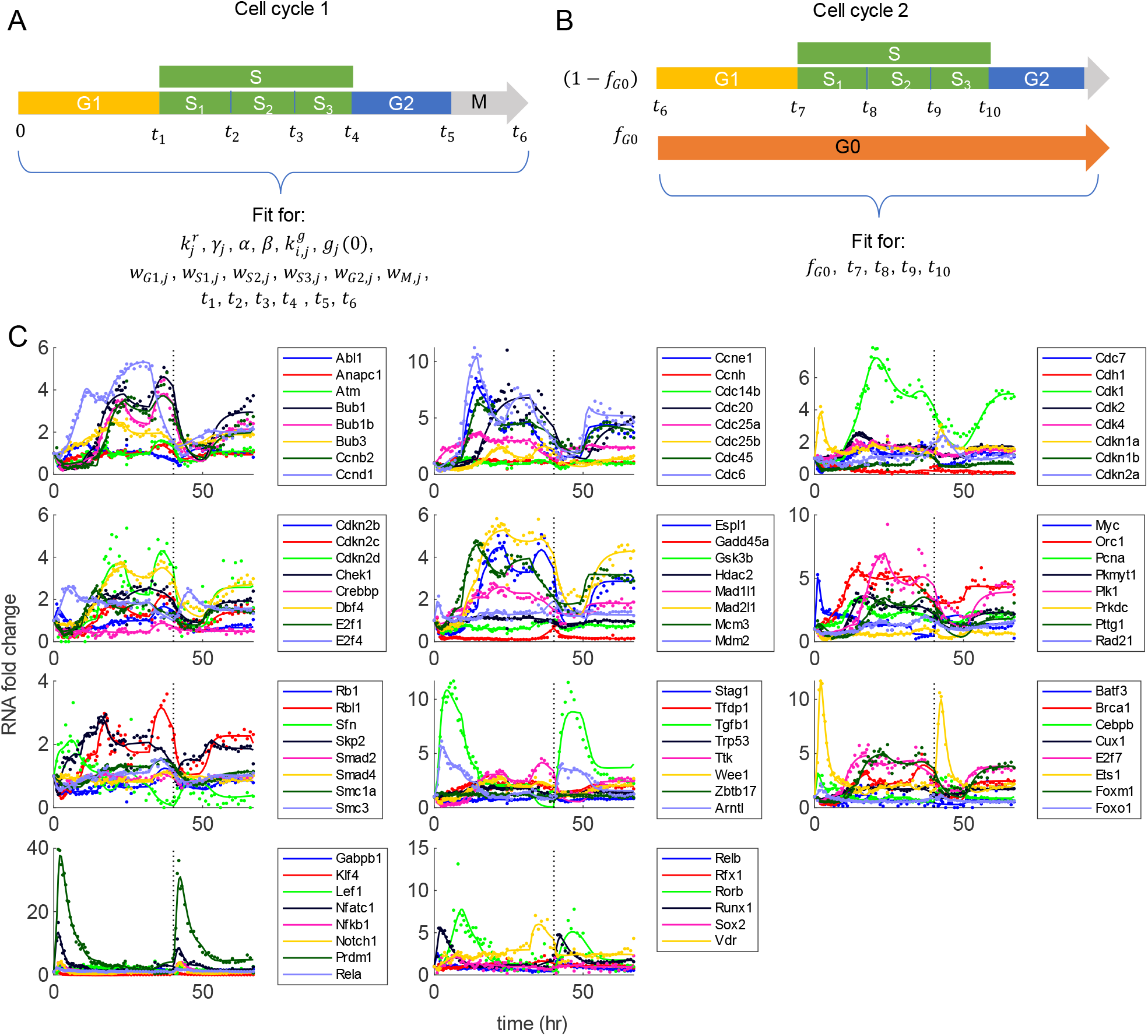
Approach for self-validating the cybernetic-inspired method (CIM). Different stages in the stage-specific cybernetic model for more than 1 cell cycle period are shown. The S phase has 3 sub-phases (represented as S1, S2, S3), with each sub-phase having its own objective. (A) During cell cycle 1, all model parameters including the RNA intrinsic parameters are fitted. (B) During cell cycle 2, fraction of cells (f_G0_) will go to cell cycle arrest (G0 phase). By only fitting for the new parameters, we validate our model. (C) Model fits to all 86 canonical cell cycle and transcription factor RNAs. The dots represent experimental data and lines represent model fits. The vertical black dotted line represents cell division.

### Model captures the experimental measurements

We applied our CIM to the selected 86 genes. Since we already identified the most likely interactions based on TDMI based time-series analysis, we can prevent the over-parameterization problem in our approach by allowing only those interactions to have a non-zero interaction parameter (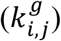) value. We had time-series measurements for 67 hours covering the period exceeding one cell cycle. After the first cell division, we assume that some cells of fraction *f*_*G*0_ could transition to G0 phase or cell cycle arrest (Figure 3A, B) while the remaining cells of fraction (1 – *f*_*G*0_) continued to the second cell cycle with objective-changing time points to be *t*_7_, *t*_8_, etc., (Figure 3A, B). We used the following strategy to validate our model. For the first cell cycle period, we fitted for all the intrinsic RNA parameters 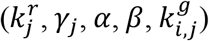, initial point for regulator (*g_j_*(0)), stage-specific weights representing their objectives (*w*_*G*1,*j*_, *w*_*s*1,*j*_, *w*_*S*2,*j*_, *w*_*S*3,*j*_, *w*_*G*2,*j*_, *w_M,j_*) and the respective time points for objective change (*t*_1_, *t*_2_, *t*_3_, *t*_4_, *t*_5_, *t*_6_). We solved the model to attain global minimization of the sum of the squares of normalized fit errors using the MATLAB ODE function “ode15s” and optimization functions “lsqnonlin” and “patternsearch” starting with 100 different initial conditions. For the time-period after the cell division, we modeled the two fractions of cells representing G0 stage and remaining cells in their second cell cycle separately and added them based on their respective fractions. Here we assumed that the G1 and G0 stages to have the same objectives and therefore equal weights (*w*_*G*0,*j*_ = *w*_*G*1,*j*_). While fitting the period after cell division, we used the same RNA intrinsic parameters and weights and fitted only for the *f*_*G*0_ and the transition times (*t*_7_, *t*_8_) (Figure 3). Figure 3C shows the overall fits for the full duration of 67 hours. Tables S1 and S2 show the overall fitted model parameters. The model described the first cell cycle and was also able to fit the time-period after cell division using the same parameters thereby providing a biologically feasible solution using our approach (this partially validates the computational model).

### Network reconstruction using CIM

Using the CIM, we reconstructed the network based on the interaction parameter 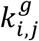. This resulted in only 153 interactions within this network of 86 nodes based on having a non-zero 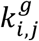 value (Figures 4A and S2). This is far less than what was predicted using the TDMI approach (731 interactions). The positive 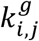 represents activation and negative 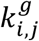 represents repression. The absolute value of 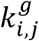 represents the strength of interaction. These interactions are identified based on a mechanistic model that is non-linear and multi-factorial, thereby making these interactions highly probable. Comparing this network with a canonical protein-protein interaction network including second neighbor interactions shows 90% of these interactions are observed in the literature (Figure S3).

**Figure 4:**
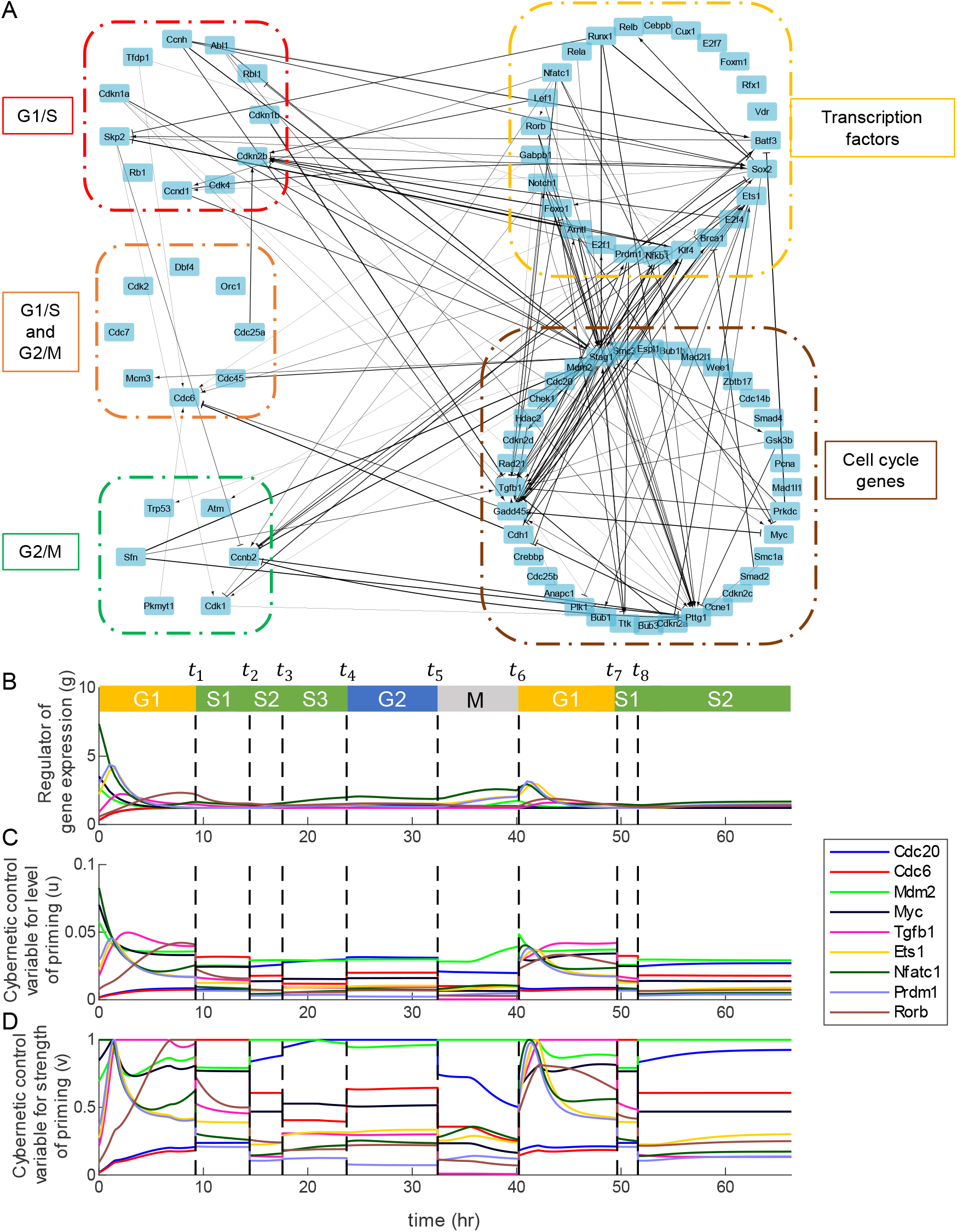
The reconstructed network based on cybernetic-inspired method (CIM) and the identification of the key regulators. (A) The activating interactions are shown as arrows and repressing interactions are shown as dashes. The thickness of the interaction lines depends on the absolute value of the interaction parameter, 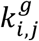. The boxed sections are color-coded based on category mentioned. The dynamics of regulator of gene expression g (B) and control variables u (C) and v (D) for the key genes. The black dashed lines represent the change points. Each region is color-coded based on its respective stage.

### Distinct stages across cell cycle and their durations

Since our approach involves changing from one stage to the next stage, it implies the involvement of change-points in our biological process and in our modeling strategy. We can use approaches such as Change-point detection (CPD) algorithms well known in signal processing community to decipher the change-points (39). We can then validate these using biological intermediate endpoints such as the those in each phase of the cell cycle. Previously, for the cell cycle system, a model-free CPD algorithm based on Singular Spectral Analysis was used to detect changes in the time series of cell cycle genes (3). The time of phase-change for the cell cycle phases was described as the time at which they identify significant individual time-series change-points detected out of 63 cell cycle genes. Based on this criterion, the duration for G1, S and G2/M phases of the cell cycle was estimated to be 14.5, 10 and 4 h, respectively (3). These CPD-algorithms are only based on time-series and lack any mechanistic insights. Here, we used the CIM to identify these change-points by determining the times at which the objective needs change. The time-points for objective change are simultaneously fitted along with other parameters. Our model predicted the duration for each of these stages {*G*1, *S*_1_, *S*_2_, *S*_3_, *G*2, *M*} during first cell cycle (Table 1). For the time after cell division, our model predicted duration for the second cycle {*G*1,*S*1} to be {9.62, 1.78} respectively. The plots of ‘g’, ‘u’ and ‘v’ of specific key regulators are shown in Figure 4B-D. The key regulators are selected based on the criterion that at least at one time point the value of ‘*v*’ peaks to 1 for them. For understanding importance of each RNA transcript, we define a new estimate called *Maximal RNA expression rate* (MRER; max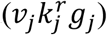). This is the maximum value among the RNA expression rates.

**Table 1.** Predicted phase durations across the first and second cell cycles. (NA – Not applicable)

| Phase duration | Cell cycle 1 (h) | Cell cycle 2 (h) |
| --- | --- | --- |
| G1 | 9.25 | 9.62 |
| S1 | 5.18 | 1.78 |
| S2 | 3.14 | >15.38 |
| S3 | 6.14 | NA |
| G2 | 8.72 | NA |
| M | 7.77 | NA |

### Network simplification and analysis

We analyze the reduced network to identify the key central molecules whose regulations are the most important for the cell cycle. We had grouped the RNA transcripts into groups such as with transcription factor genes, check-point genes (G1/S and G2/M) and other canonical cell cycle genes (Figures 2G and 4A). For identifying key central nodes, we used the following two properties. One is a network property called *Maximal Clique Centrality* (MCC) that analyzes and identifies the key nodes giving importance to number and extent of interactions (40). The top nodes based on MCC were Stag1, Tgfb1 and Pttg1. Another property from the CIM is called MRER (see previous section) that helps identify key nodes in a specific-rate manner. Using thresholds for these properties, we were able to eliminate the nodes in the network while retaining the important nodes. We illustrate a simplified network based on the following thresholds (MCC≥7 ∪ MRER>1) (Figure 5). This approach reduced the nodes to 66 (from 86) and interactions to 126 (from 153). We further show parts of this simplified network by grouping them into cell cycle genes (Figure 6), transcription factor genes (Figure 7A), and check-point genes (Figure 7B, C and D).

**Figure 5:**
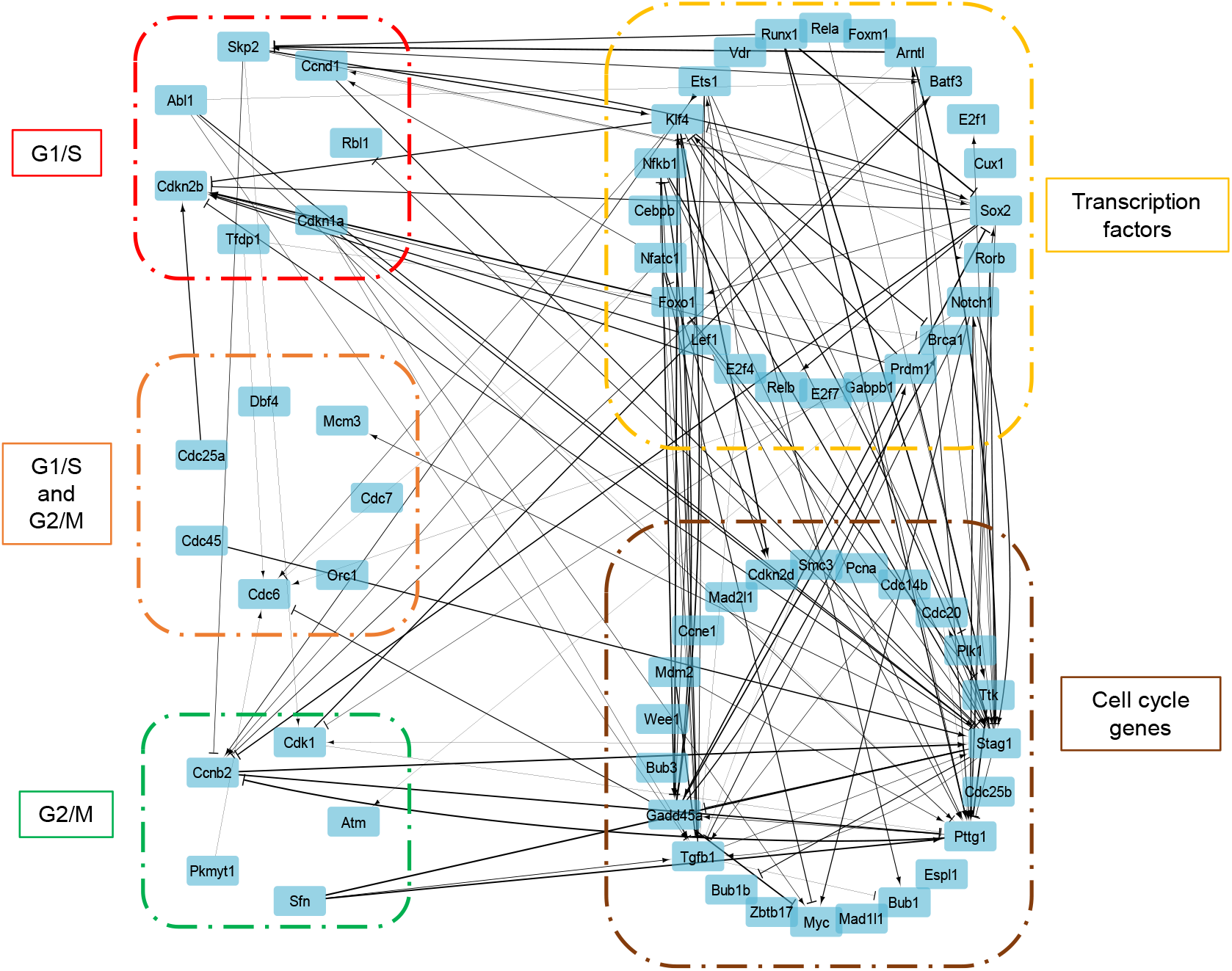
Development of simplified network based on Maximal Clique Centrality (MCC) and Maximal RNA expression rate (MRER). The activating interactions are shown as arrows and repressing interactions are shown as dashes. The thickness of the interaction lines depends on the absolute value of the interaction parameter, 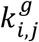. The boxed sections are color-coded based on category mentioned. The network shown here is for a union of nodes with MCC≥ 7 and MRER>1. The boxed sections are color-coded based on the category mentioned.

**Figure 6:**
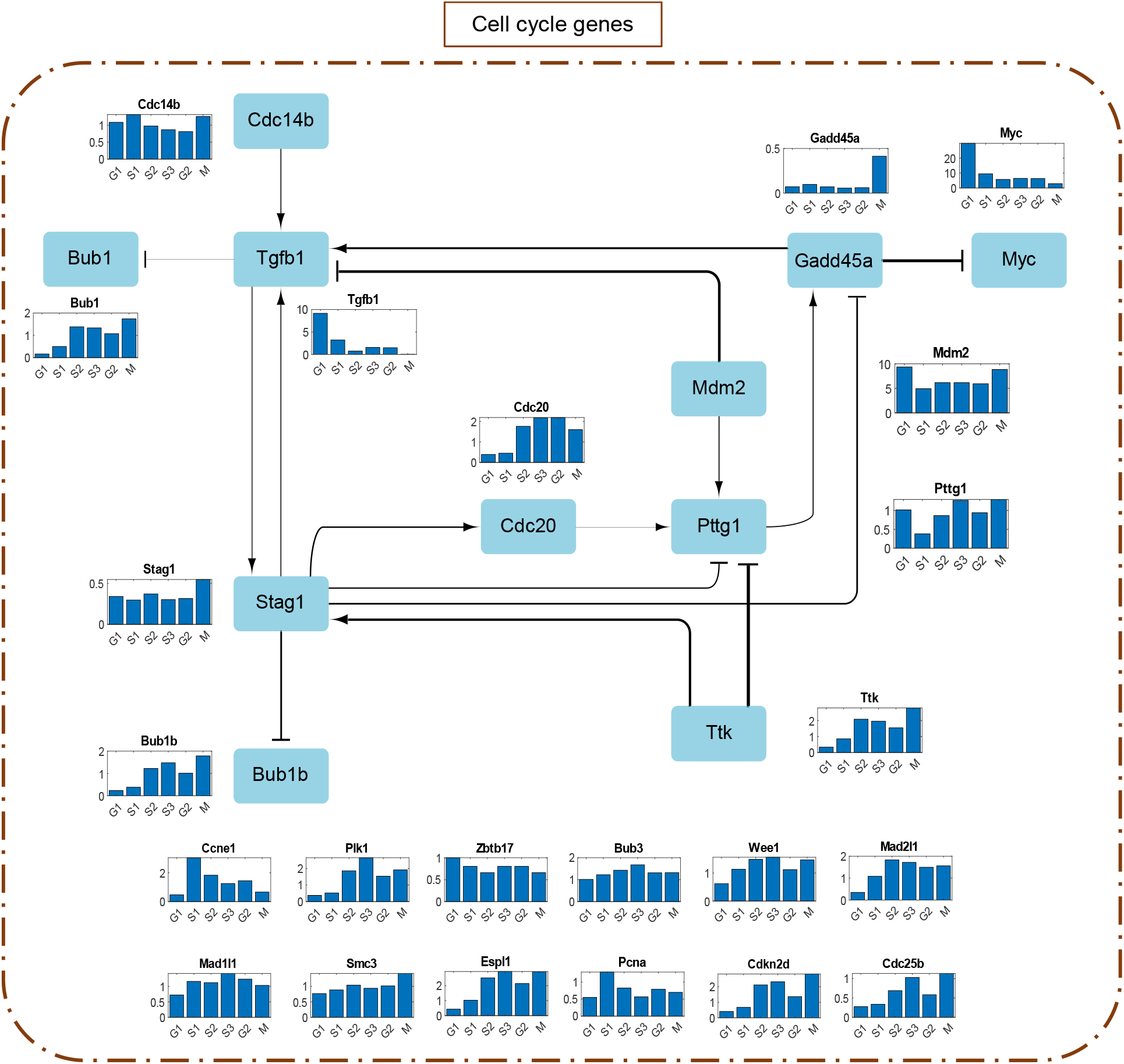
Simplified network involving only canonical cell cycle genes without the checkpoint genes and their interactions. The activating interactions are shown as arrows and repressing interactions are shown as dashes. The thickness of the interaction lines depends on the absolute value of the interaction parameter, 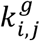. The ones without cross-interactions within the group are shown below the network. The bar plots represent the stage-specific maximal RNA expression rate (MRER).

**Figure 7:**
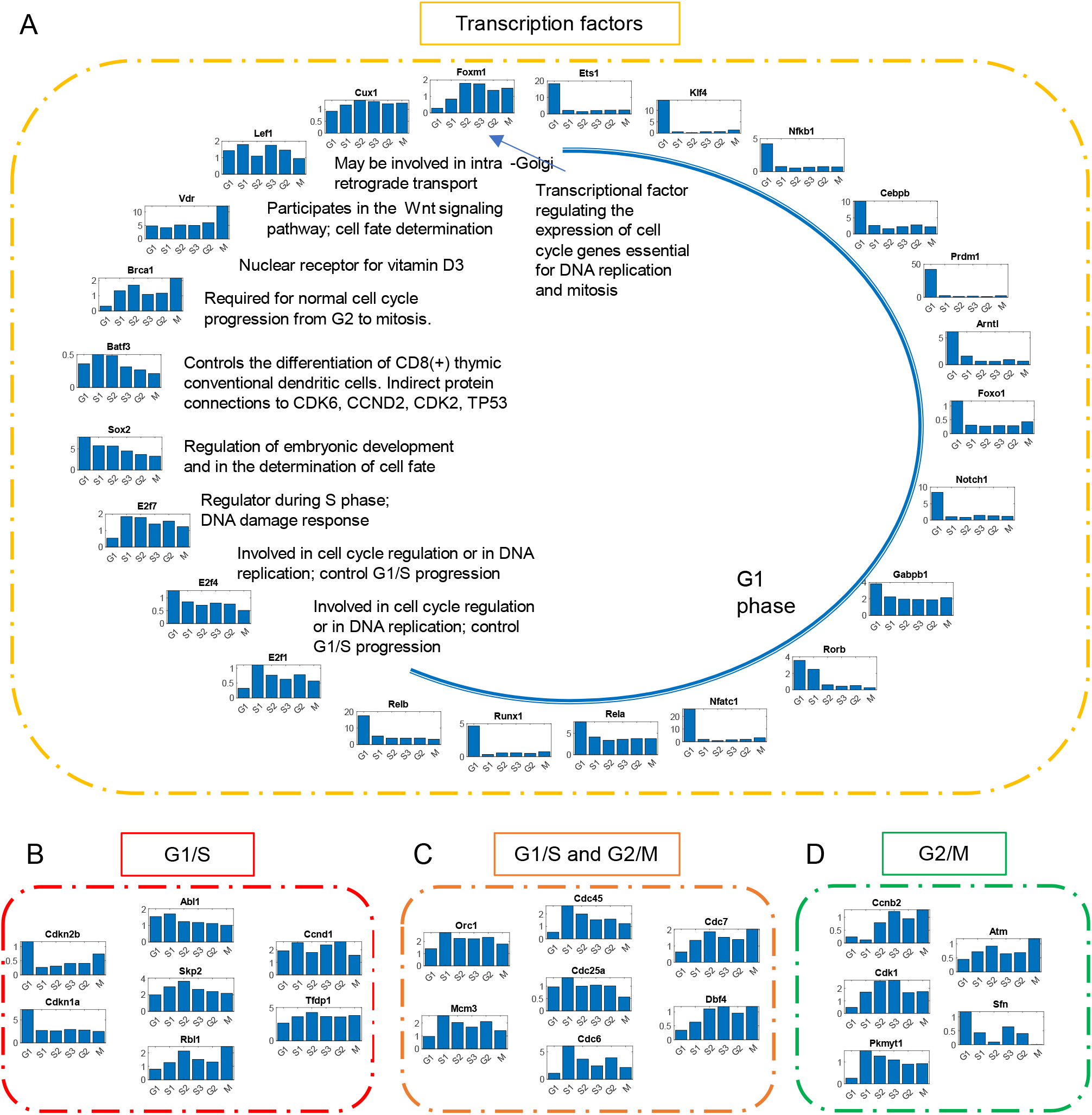
Nodes of the simplified network of (A) Transcription factor genes, (B) G1/S checkpoint genes, (C) Genes considered both G1/S and G2/M checkpoints, and (D) G2/M checkpoint genes. The bar plots represent the stage-specific maximal RNA expression rate (MRER).

## DISCUSSION

Cybernetic methods were originally developed to address unknown regulatory events in a single stage cellular process with a defined goal. However, several biological systems like the mammalian cell cycle involve multiple stages with distinct one or more objective functions at each stage. The traditional cybernetic approach is not designed to capture this complexity. Here, we develop a novel strategy for capturing this complexity, and we call it a cybernetic-inspired method (CIM). In order to account for each stage and how the stage proceeds in terms of a causal evolution, we introduced TDMI which uses temporal correlation to provide insights into the causality of regulatory events. The preliminary regulatory network from TDMI is then used as a starting point in the CIM to obtain biologically realistic models. Such pre-processing techniques have greater importance when solving inverse biological problems. The CIM approach significantly reduced the number of interactions to describe the system (compare Figure 2G and Figure 4A). While TDMI is one way to identify causality and determine concomitant interactions, we could have used other techniques for time-series analysis such as Granger causality to identify these interactions. Knowledge embedded in databases such as KEGG pathways (41), Reactome pathway database (42) and ConsensusPathDB (43) can be used for a comprehensive list of legacy pathways to ensure the validity of the initial network. While extracting an initial network directly from such databases are possible, our approach using TDMI (or other time-series analysis techniques) can better formulate this initial network incorporating causal connections when large longitudinal data are available.

Since we are introducing objective functions at distinct stages, we can use stages as defined by experimental measurements if available. However, given the dynamical nature of the system, we may have changes of state which are distinct from the experimentally determined stages. This motivated us to use CIM for change-point detection which provides insights into stages of the system, in addition to providing an opportunity to introduce novel objective functions at each of these intermediate stages. We have already shown that the S phase shows differential regulation because of the balance between transcription and replication events and requires multiple stage-changes to model them. Since we are modeling regulation within the transcription process, our stage-changes are expected to occur before we phenotypically observe them, and this time-delay can be attributed to lag between RNA expression and protein production. The dynamical evolution for the stages provides insights into the regulatory mechanisms associated with the causality leading to the global end point of the biological system in this case, cell cycle.

We observe that the stage-durations during the second cell cycle are not same as the first. In fact, during the second cell cycle, we did not identify any change-point after 51.61 hours (representing *S_2_* stage) implying that the cell cycle has slowed down or stopped. The reason for cell cycle arrest could be multifaceted with one of them being attributed to *Cdc6*. *Cdc6*, a gene involved in initiating DNA replication, is low during the second cell cycle when compared with the first cell cycle, and this could contribute to stopping the cell cycle. For a cell “cycle” model, our approach should have an ability to show a cyclic behavior if the objectives are repeated. Towards testing whether the model will yield cyclic solutions for the cell cycle, we repeated the pattern of objectives based on the first cell cycle and this resulted in a cyclic behavior (Figure S4). Each of these stage objectives are represented by their weights (*w*_*G*1,*j*_, *w*_*s*1,*j*_, *w*_*S*2,*j*_, *w*_*S*3,*j*_, *w*_*G*2,*j*_, *w_M,j_*) Figure S5). When comparing for weight difference across adjacent stages (Figure S6), we observed that the stages *S*_2_, *S*_3_, *G*2 had highly similar objectives. It is during these stages that the cell proceeds to division and having similar objectives is interesting and warrants further studies.

The CIM approach has several quantitative aspects that can be used to infer key regulators in multiple ways. One way is by using the stage-specific weights that provided us with the quantitative relevance of key players within each stage. From the estimated weights (Figure S5), *Cdc20* is important during the *S*_2_, *S*_3_, *G*2 stages as its protein is required for nuclear movement prior to anaphase and chromosome separation. We can also identify key regulators from the regulator of gene expression and the control variables (Figure 4B-D). For *G*1 phase; *Tgfb1*, *Nfatc1*, *Myc*, *Ets1*, *Rorb* and *Prdm1* are key regulators (Figure 4B-D). Tgfb1 protein is a multifunctional protein that controls proliferation, differentiation and other functions in many cell types and is known to participate in regulation of the G1/S checkpoint. For *S*_1_ phase, *Cdc6* is a key regulator as its protein is known to be involved in the initiation of DNA replication and also participates in checkpoint controls that ensure DNA replication is completed before mitosis is initiated (Figure 4B-D). During *S*_2_, *S*_3_ and *M* phases, *Mdm2* is a key regulator since its protein is known to inhibit p53/TP53- and p73/TP73-mediated cell cycle arrest and thereby preventing cell cycle arrest. Here, *Cdc20* is a key regulator for *S*_3_ and *G*2 phases. For understanding phase-specific importance for each RNA transcript, we used MRER (Figure S7).

We further simplified the network based on a combination of network and dynamical property (MCC and MRER, respectively). Figure 6 shows the part of simplified network involving only canonical cell cycle genes without the check-point genes. This network provides us with players required for the cell cycle. *Tgfb1*, *Stag1* and *Pttg1* are key regulators with high degrees. *Tgfb1* has high MRER during *G*1 phase representing its key participation in the regulation of G1/S checkpoint. Stag1 has high MRER during *M* phase as they are required in the cohesion of sister chromatids after DNA replication. *Pttg1* has high MRER during *S*_3_ and *M* phases because of its central role in chromosome stability, p53/TP53 pathway, and DNA repair. *Gadd45a*, known to stimulate DNA excision repair in vitro and inhibit entry of cells into S phase, negatively regulates Myc that activates transcription of growth-related proteins. Most of these RNAs have high MRER during the later stages of cell cycle since multiple players are required during division. Figure 7 shows the nodes of the simplified network of transcription factor genes and checkpoint genes. Unlike canonical cell cycle genes, most of the transcription factor genes have high MRER during the *G1* phase and possibly because of high requirement of transcription factors after cell division for cellular growth. Other transcription factor genes have some role in the cell cycle as shown in Figure 7A. *Vdr*, a nuclear receptor for vitamin D3 also supports the cellular growth (44). *Batf3* can also support the cell cycle based on protein-protein interaction with CDK6, CCND2, CDK2, and TP53. The checkpoint genes show high variability based on phase-specific MRER values (Figures 7 B, C, and D) showing that they are expressed at various timepoints but can act as phase-specific checkpoints.

While there were several attempts to develop transcriptional regulatory networks (45, 46), these approaches were not extended to dynamically model longitudinal measurements. Also, these eukaryotic systems show complex regulatory patterns which simple linear models cannot capture. The ability of the cybernetic approach to address this regulatory complexity lies in the nonlinearity enforced by the cybernetic control variables as well as the segregation of the system into multiple stages with varying control objectives. In our approach, we have used ‘time’ to characterize all transitions because of its inherent simplicity. However, we recognize that such transitions may be related more deeply to intracellular variables, a potential that can only be realized when sufficient understanding of the system becomes available. As we have modeled a synchronous cell cycle here all cells may be assumed to behave similarly. On the other hand, most biological experiments are asynchronous, necessitating some form of averaging to be superimposed on the model where we can model cell behavior based on averaging weights over the stage-specific number density of cells.

The complexity within cellular processes due to multitude of regulatory interactions is often difficult to infer using previous mathematical models. The cybernetic approach of mapping the cellular regulation to objectives that can be mathematically formulated can overcome this difficulty. Based on the observation that the cellular processes are multi-staged, we developed a novel approach that incorporates the stage-specific objectives. The implications of our proposed approach go beyond modeling the mammalian cell cycle processes. We can formulate complex cellular processes, where only sparse measurements are available, albeit with knowledge of intermediate end points, in terms of the CIM, enabling us to infer multiple unknown regulatory processes. For instance, we can consider a developmental process of lineage specification of a specialized tissue from pluripotent stem cells, where targeted measurements are available across stages of development (pseudo time) as an exemplary problem for this approach. While such investigations are in progress, it must be evident that the cybernetic approach by comprehensive addition of regulatory intervention, has a higher potential to discover new biological phenomena.

## AUTHOR CONTRIBUTIONS

R.R. wrote the manuscript; M.R.M., S.S. and D. R. revised the manuscript; R.R., S.S. and D. R. designed the research; R.R. performed the research; R.R. analyzed the data; S.K., L.A., M.R.M., S.G., S.S. and D.R. contributed analytical tools.

## ACKNOWLEDGEMENTS

This work was supported by NIH GRANTS, R01 LM012595 (S.S.), OT2 OD030544 (S.S.), R01 HL106579 (S.S.), R01 HL108735 (S.S.), U2C DK119886 (S.S.), The Joan And Irwin Jacobs Endowed Professorship (S.S.), Center For Science Of Information (CSOI), A National Science Foundation Science And Technology Center, under Grant Agreement CCF-0939370 (S.S. AND D.R.) AND The Harry Creighton Peffer Endowed Professorship (D.R.).

## SUPPORTING INFORMATION LEGENDS

Figure S1: Development of interaction network using TDMI values. The heatmap shows the maximum TDMI value between two RNAs across different delays. Only interactions with TDMI of top 0.1 of the TDMI values are shown.

Figure S2: Heat map for the interaction parameter 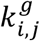. Red color represents activation interaction and blue color represents repressive interaction.

Figure S3: Comparing our network with String database. Comparing the network with a canonical protein-protein interaction network including second neighbor interactions shows 90% of these interactions are validated.

Figure S4: Cyclicity in our model. The figure shows that if we repeat the objectives for each phase in the same order, the model shows a cyclic behavior.

Figure S5: Stage-specific cybernetic weights.

Figure S6: Cybernetic weight differences between adjacent stages.

Figure S7: Stage-specific Maximal RNA expression rate (Max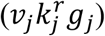))

Table S1: Cybernetic model RNA species-agnostic parameters

Table S2: Cybernetic model RNA species-specific parameters

