## Supporting Information for "Modeling transcriptional regulation of the cell cycle using a novel cybernetic-inspired approach"

*Table S1: Cybernetic model RNA species-agnostic parameters*

| Parameters | Value | Unit |
|---|---|---|
| $\alpha$ | 0.7705 | g-unit/h |
| $\beta$ | 0.6349 | $h^{-1}$ |
| $f_{G0}$ | 0.3091 | - |
| $t_1$ | 9.2511 | h |
| $t_2$ | 14.4324 | h |
| $t_3$ | 17.5751 | h |
| $t_4$ | 23.7185 | h |
| $t_5$ | 32.4408 | h |
| $t_6$ | 40.2104 | h |
| $t_7$ | 49.8328 | h |
| $t_8$ | 51.6155 | h |

Here, 'g-unit' is the basic unit of the regulator of gene expression (*g*).

*Table S2: Cybernetic model RNA species-specific parameters*

| Parameter | $k_j^r$ | $g_j(0)$ | $\gamma_j$ | $w_{G1,j}$ | $w_{S1,j}$ | $w_{S2,j}$ | $w_{S3,j}$ | $w_{G2,j}$ | $w_{M,j}$ |
|---|---|---|---|---|---|---|---|---|---|
| Unit | $h^{-1}$ g-unit$^{-1}$ | g-unit | $h^{-1}$ | - | - | - | - | - | - |
| Abl1 | 2.070 | 0.920 | 1.036 | 0.016 | 0.014 | 0.014 | 0.012 | 0.012 | 0.010 |
| Anapc1 | 1.701 | 1.070 | 0.654 | 0.013 | 0.009 | 0.011 | 0.012 | 0.012 | 0.016 |
| Atm | 1.895 | 0.378 | 0.699 | 0.011 | 0.013 | 0.014 | 0.010 | 0.011 | 0.017 |
| Bub1 | 1.914 | 0.316 | 0.337 | 0.004 | 0.009 | 0.021 | 0.020 | 0.017 | 0.024 |
| Bub1b | 2.163 | 0.316 | 0.424 | 0.005 | 0.005 | 0.014 | 0.017 | 0.013 | 0.020 |
| Bub3 | 2.581 | 1.283 | 0.675 | 0.014 | 0.012 | 0.012 | 0.014 | 0.012 | 0.010 |
| Ccnb2 | 2.009 | 0.447 | 0.318 | 0.004 | 0.002 | 0.011 | 0.017 | 0.014 | 0.016 |
| Ccnd1 | 1.869 | 1.781 | 0.489 | 0.015 | 0.014 | 0.013 | 0.020 | 0.023 | 0.013 |
| Ccne1 | 2.689 | 0.316 | 0.316 | 0.006 | 0.026 | 0.014 | 0.010 | 0.012 | 0.005 |
| Ccnh | 1.487 | 1.251 | 0.343 | 0.015 | 0.011 | 0.009 | 0.009 | 0.011 | 0.010 |
| Cdc14b | 2.140 | 1.525 | 0.888 | 0.010 | 0.008 | 0.008 | 0.009 | 0.009 | 0.012 |
| Cdc20 | 1.537 | 0.317 | 0.316 | 0.015 | 0.012 | 0.036 | 0.038 | 0.041 | 0.026 |
| Cdc25a | 1.664 | 2.039 | 0.331 | 0.015 | 0.016 | 0.011 | 0.011 | 0.011 | 0.005 |
| Cdc25b | 1.658 | 0.317 | 0.412 | 0.009 | 0.008 | 0.014 | 0.021 | 0.013 | 0.021 |
| Cdc45 | 1.538 | 0.329 | 0.316 | 0.013 | 0.024 | 0.016 | 0.014 | 0.017 | 0.011 |
| Cdc6 | 4.973 | 0.316 | 0.547 | 0.004 | 0.015 | 0.008 | 0.005 | 0.009 | 0.004 |
| Cdc7 | 2.267 | 0.317 | 1.111 | 0.011 | 0.017 | 0.020 | 0.017 | 0.016 | 0.021 |
| Cdh1 | 0.768 | 1.842 | 0.464 | 0.017 | 0.015 | 0.005 | 0.007 | 0.007 | 0.011 |
| Cdk1 | 1.740 | 0.316 | 0.316 | 0.012 | 0.015 | 0.014 | 0.014 | 0.015 | 0.015 |
| Cdk2 | 1.819 | 1.789 | 0.316 | 0.010 | 0.017 | 0.011 | 0.009 | 0.010 | 0.009 |
| Cdk4 | 1.560 | 1.697 | 0.316 | 0.015 | 0.017 | 0.012 | 0.013 | 0.013 | 0.010 |
| Cdkn1a | 3.623 | 2.574 | 1.647 | 0.015 | 0.008 | 0.006 | 0.007 | 0.007 | 0.005 |
| Cdkn1b | 1.451 | 0.316 | 0.554 | 0.011 | 0.006 | 0.011 | 0.009 | 0.009 | 0.018 |
| Cdkn2a | 1.312 | 1.014 | 0.316 | 0.011 | 0.009 | 0.011 | 0.012 | 0.013 | 0.010 |
| Cdkn2b | 1.440 | 4.966 | 0.403 | 0.008 | 0.008 | 0.008 | 0.011 | 0.012 | 0.018 |
| Cdkn2c | 1.511 | 0.319 | 0.363 | 0.006 | 0.011 | 0.017 | 0.015 | 0.011 | 0.020 |
| Cdkn2d | 2.403 | 0.317 | 0.615 | 0.006 | 0.007 | 0.020 | 0.022 | 0.014 | 0.025 |
| Chek1 | 1.610 | 0.430 | 0.323 | 0.009 | 0.017 | 0.017 | 0.014 | 0.019 | 0.015 |
| Crebbp | 1.548 | 0.609 | 0.584 | 0.012 | 0.006 | 0.007 | 0.006 | 0.007 | 0.012 |
| Dbf4 | 1.797 | 1.139 | 0.319 | 0.010 | 0.012 | 0.019 | 0.020 | 0.018 | 0.019 |
| E2f1 | 1.800 | 0.316 | 0.417 | 0.009 | 0.021 | 0.013 | 0.011 | 0.014 | 0.009 |
| E2f4 | 1.701 | 2.419 | 0.407 | 0.013 | 0.012 | 0.009 | 0.009 | 0.009 | 0.005 |
| Espl1 | 2.573 | 0.317 | 0.650 | 0.006 | 0.010 | 0.021 | 0.024 | 0.019 | 0.023 |
| Gadd45a | 0.381 | 1.119 | 0.469 | 0.007 | 0.011 | 0.016 | 0.013 | 0.018 | 0.021 |
| Gsk3b | 1.174 | 0.495 | 0.320 | 0.010 | 0.007 | 0.008 | 0.007 | 0.009 | 0.014 |
| Hdac2 | 1.694 | 1.443 | 0.460 | 0.012 | 0.010 | 0.010 | 0.009 | 0.011 | 0.009 |
| Mad1l1 | 1.968 | 1.148 | 0.538 | 0.017 | 0.019 | 0.016 | 0.020 | 0.019 | 0.014 |
| Mad2l1 | 2.065 | 0.318 | 0.316 | 0.007 | 0.016 | 0.024 | 0.022 | 0.021 | 0.019 |
| Mcm3 | 2.790 | 0.317 | 0.529 | 0.011 | 0.021 | 0.014 | 0.012 | 0.016 | 0.009 |
| Mdm2 | 5.074 | 2.653 | 4.521 | 0.019 | 0.012 | 0.013 | 0.013 | 0.014 | 0.012 |
| Myc | 10.011 | 3.515 | 4.803 | 0.009 | 0.006 | 0.003 | 0.004 | 0.004 | 0.001 |
| Orc1 | 2.559 | 0.368 | 0.453 | 0.020 | 0.026 | 0.019 | 0.019 | 0.021 | 0.014 |

| | | | | | | | | | |
|---|---|---|---|---|---|---|---|---|---|
| Pcna | 1.886 | 1.928 | 0.325 | 0.013 | 0.023 | 0.013 | 0.009 | 0.013 | 0.010 |
| Pkmyt1 | 1.582 | 0.316 | 0.317 | 0.009 | 0.020 | 0.015 | 0.015 | 0.015 | 0.013 |
| Plk1 | 2.185 | 0.319 | 0.354 | 0.007 | 0.007 | 0.021 | 0.031 | 0.019 | 0.021 |
| Prkdc | 1.186 | 0.318 | 0.471 | 0.007 | 0.008 | 0.011 | 0.010 | 0.013 | 0.020 |
| Pttg1 | 1.698 | 7.375 | 0.339 | 0.002 | 0.008 | 0.016 | 0.024 | 0.019 | 0.023 |
| Rad21 | 1.350 | 0.621 | 0.316 | 0.013 | 0.009 | 0.016 | 0.014 | 0.013 | 0.019 |
| Rb1 | 1.264 | 0.525 | 0.317 | 0.010 | 0.007 | 0.011 | 0.007 | 0.008 | 0.015 |
| Rbl1 | 2.506 | 0.317 | 0.750 | 0.011 | 0.013 | 0.019 | 0.013 | 0.012 | 0.020 |
| Sfn | 0.644 | 6.667 | 0.321 | 0.024 | 0.031 | 0.008 | 0.011 | 0.007 | 0.000 |
| Skp2 | 2.932 | 0.316 | 1.161 | 0.013 | 0.014 | 0.015 | 0.015 | 0.017 | 0.013 |
| Smad2 | 1.596 | 1.336 | 0.366 | 0.011 | 0.010 | 0.008 | 0.007 | 0.008 | 0.008 |
| Smad4 | 1.629 | 1.904 | 0.317 | 0.011 | 0.006 | 0.005 | 0.006 | 0.006 | 0.008 |
| Smc1a | 1.435 | 1.810 | 0.318 | 0.009 | 0.010 | 0.010 | 0.010 | 0.010 | 0.011 |
| Smc3 | 2.335 | 1.235 | 0.911 | 0.013 | 0.010 | 0.010 | 0.009 | 0.011 | 0.013 |
| Stag1 | 1.414 | 0.559 | 0.380 | 0.009 | 0.006 | 0.008 | 0.008 | 0.008 | 0.011 |
| Tfdp1 | 3.729 | 1.025 | 1.674 | 0.017 | 0.015 | 0.015 | 0.014 | 0.015 | 0.013 |
| Tgfb1 | 4.087 | 0.903 | 0.703 | 0.021 | 0.008 | 0.002 | 0.005 | 0.005 | 0.000 |
| Trp53 | 1.989 | 1.762 | 0.326 | 0.010 | 0.011 | 0.007 | 0.008 | 0.009 | 0.005 |
| Ttk | 2.397 | 0.316 | 0.634 | 0.005 | 0.009 | 0.020 | 0.019 | 0.016 | 0.025 |
| Wee1 | 2.458 | 0.771 | 0.594 | 0.009 | 0.012 | 0.013 | 0.014 | 0.011 | 0.012 |
| Zbtb17 | 2.375 | 2.232 | 0.643 | 0.013 | 0.009 | 0.006 | 0.008 | 0.008 | 0.006 |
| Arntl | 3.121 | 1.192 | 0.691 | 0.021 | 0.010 | 0.004 | 0.004 | 0.006 | 0.003 |
| Batf3 | 1.349 | 3.658 | 0.396 | 0.005 | 0.012 | 0.010 | 0.007 | 0.009 | 0.006 |
| Brca1 | 2.260 | 0.317 | 0.511 | 0.006 | 0.016 | 0.018 | 0.012 | 0.014 | 0.022 |
| Cebpb | 6.628 | 2.775 | 2.950 | 0.011 | 0.004 | 0.002 | 0.003 | 0.004 | 0.003 |
| Cux1 | 2.357 | 1.027 | 0.714 | 0.015 | 0.013 | 0.014 | 0.013 | 0.013 | 0.012 |
| E2f7 | 2.143 | 0.316 | 0.361 | 0.011 | 0.025 | 0.021 | 0.017 | 0.020 | 0.014 |
| Ets1 | 4.454 | 2.226 | 1.096 | 0.010 | 0.007 | 0.003 | 0.004 | 0.004 | 0.003 |
| Foxm1 | 2.006 | 0.323 | 0.344 | 0.006 | 0.013 | 0.024 | 0.024 | 0.020 | 0.019 |
| Foxo1 | 1.306 | 3.130 | 0.492 | 0.015 | 0.011 | 0.009 | 0.010 | 0.008 | 0.007 |
| Gabpb1 | 3.197 | 3.034 | 1.140 | 0.015 | 0.009 | 0.006 | 0.006 | 0.006 | 0.006 |
| Klf4 | 2.238 | 9.998 | 5.455 | 0.011 | 0.005 | 0.002 | 0.002 | 0.002 | 0.003 |
| Lef1 | 1.960 | 2.579 | 0.598 | 0.014 | 0.015 | 0.008 | 0.009 | 0.008 | 0.005 |
| Nfatc1 | 3.504 | 7.366 | 0.957 | 0.014 | 0.005 | 0.002 | 0.003 | 0.003 | 0.003 |
| Nfkb1 | 2.596 | 2.767 | 0.819 | 0.012 | 0.006 | 0.003 | 0.004 | 0.004 | 0.003 |
| Notch1 | 2.197 | 3.132 | 1.693 | 0.012 | 0.008 | 0.004 | 0.004 | 0.004 | 0.003 |
| Prdm1 | 9.708 | 3.003 | 0.576 | 0.005 | 0.002 | 0.001 | 0.001 | 0.001 | 0.001 |
| Rela | 6.312 | 2.780 | 3.244 | 0.009 | 0.005 | 0.004 | 0.004 | 0.004 | 0.003 |
| Relb | 6.405 | 3.352 | 5.961 | 0.014 | 0.008 | 0.005 | 0.005 | 0.006 | 0.004 |
| Rfx1 | 1.791 | 1.358 | 0.423 | 0.012 | 0.009 | 0.009 | 0.010 | 0.010 | 0.012 |
| Rorb | 1.536 | 0.575 | 0.377 | 0.039 | 0.020 | 0.009 | 0.007 | 0.008 | 0.004 |
| Runx1 | 1.710 | 4.030 | 0.435 | 0.020 | 0.007 | 0.011 | 0.007 | 0.006 | 0.005 |
| Sox2 | 4.732 | 0.576 | 3.680 | 0.017 | 0.009 | 0.008 | 0.010 | 0.010 | 0.007 |
| Vdr | 10.034 | 0.743 | 2.026 | 0.004 | 0.003 | 0.003 | 0.003 | 0.004 | 0.006 |

Here, 'g-unit' is the basic unit of the regulator of gene expression (*g*).

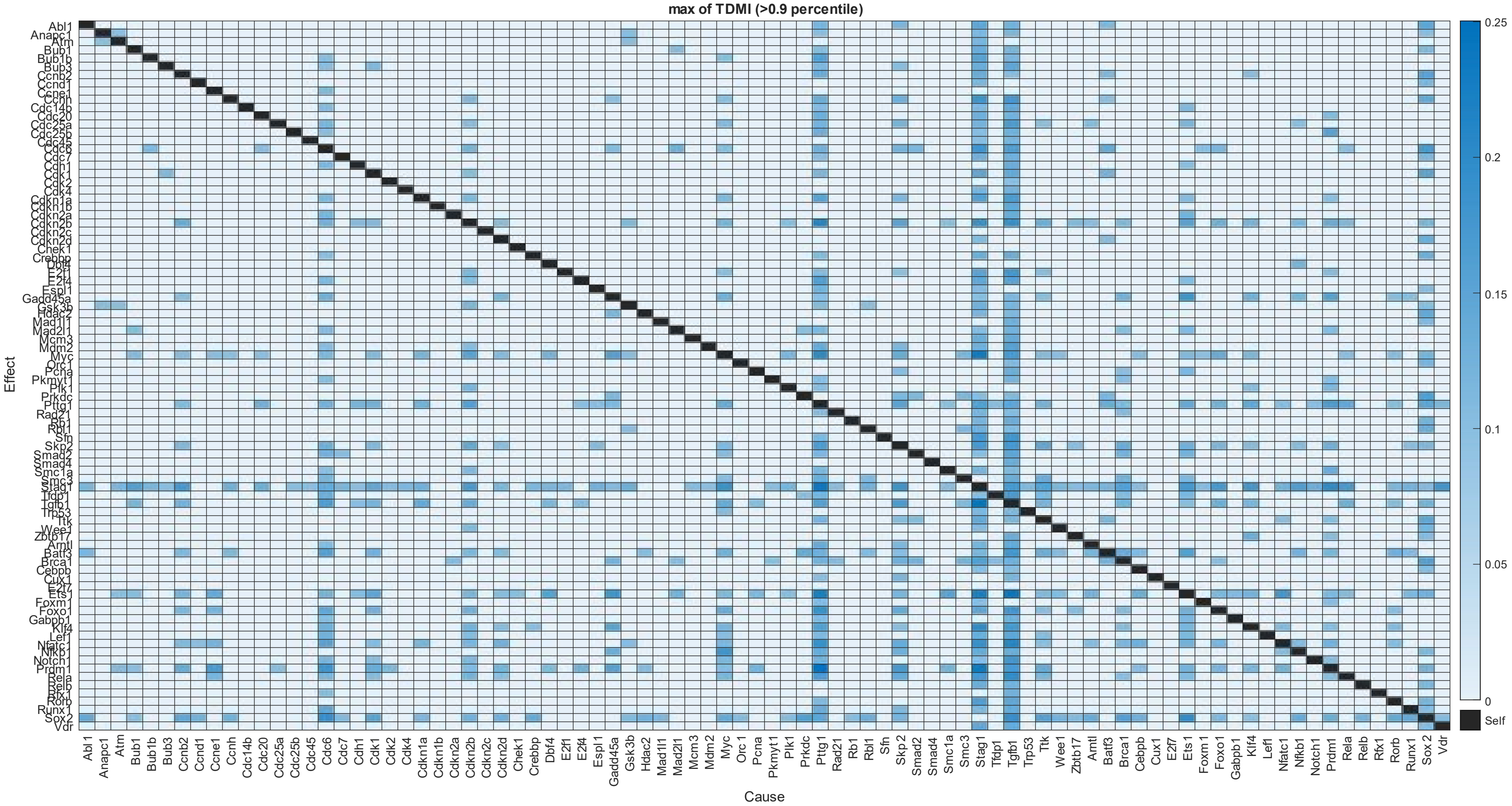


*Figure S1: Development of interaction network using TDMI values. The heatmap shows the maximum TDMI value between two RNAs across different delays. Only interactions with TDMI of top 0.1 of the TDMI values are shown.*

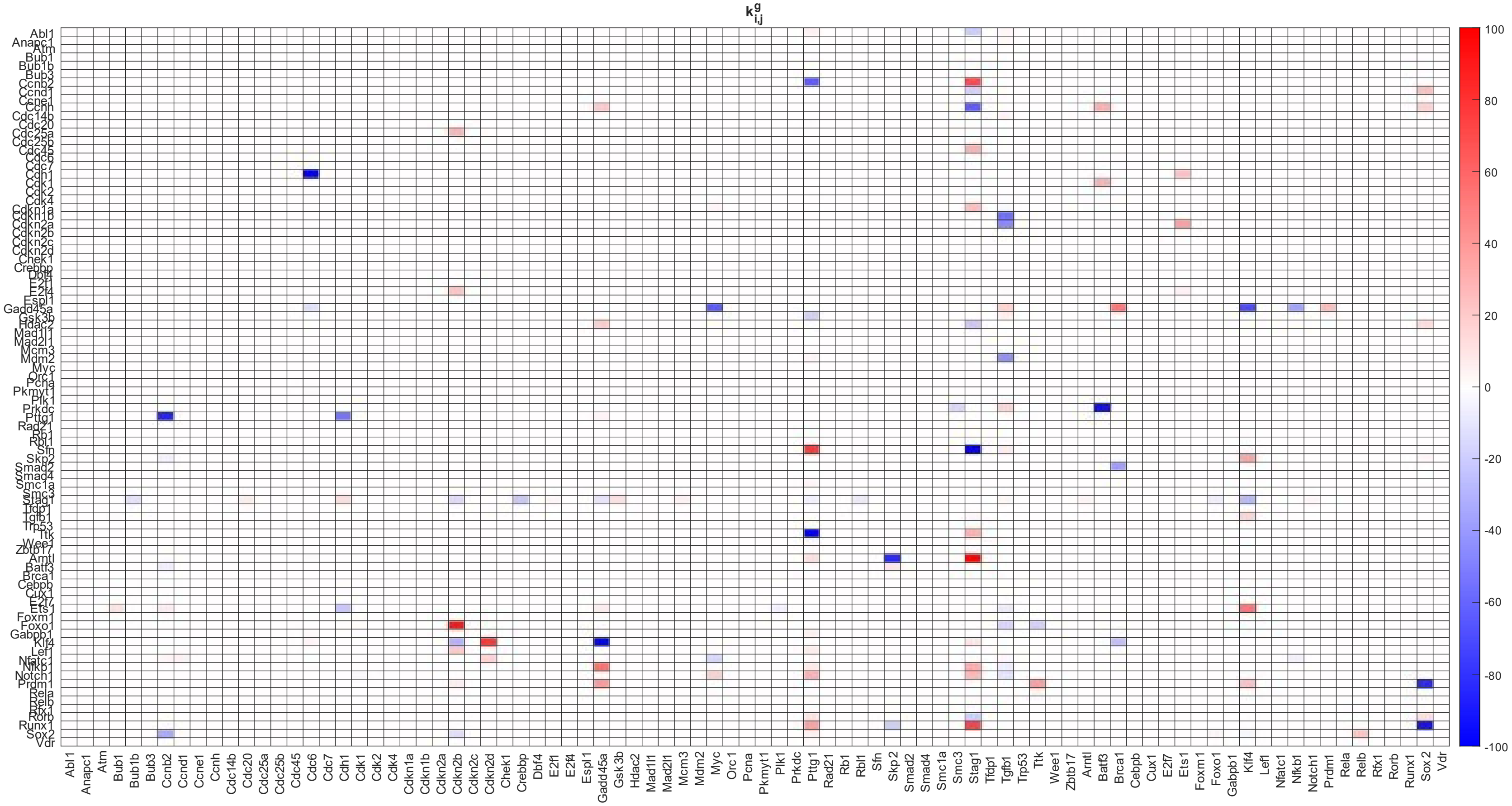


*Figure S2: Heat map for the interaction parameter $k^g_{i,j}$. Red color represents activation interaction and blue color represents repressive interaction.*

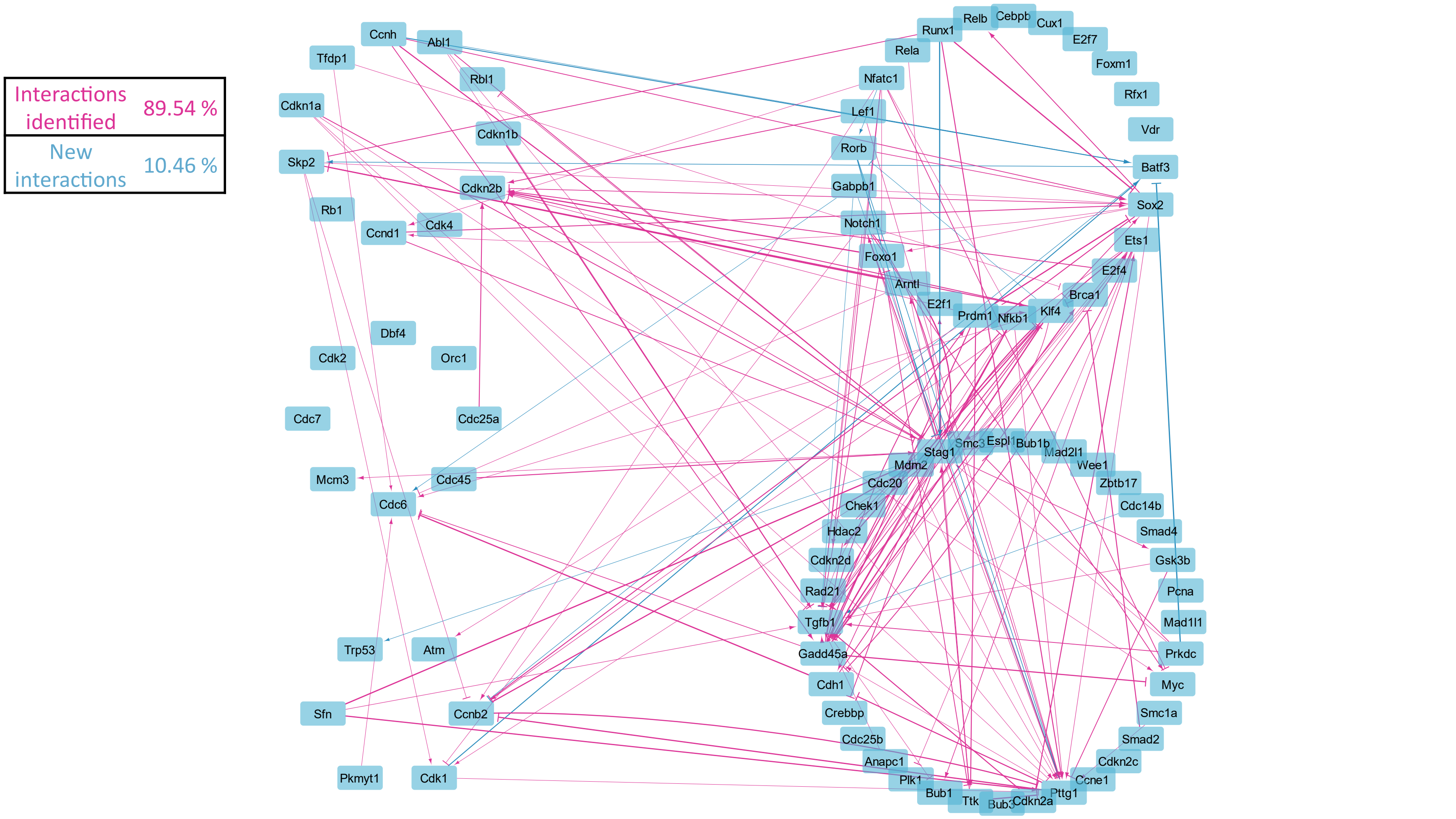


*Figure S3: Comparing our network with String database. Comparing the network with a canonical protein-protein interaction network including second neighbor interactions shows 90% of these interactions are validated.*

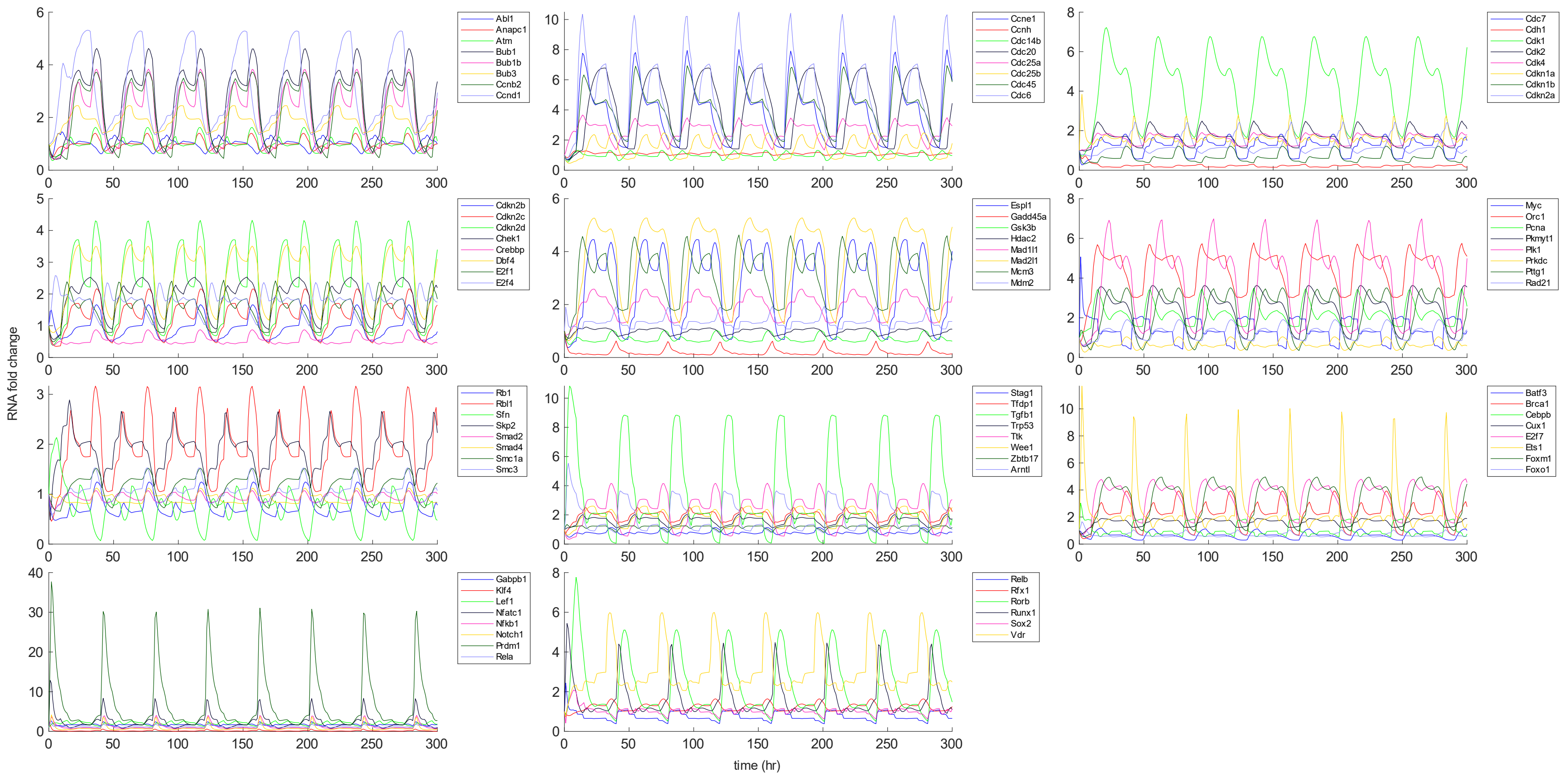


*Figure S4: Cyclicity in our model. The figure shows that if we repeat the objectives for each phase in the same order, the model shows a cyclic behavior.*

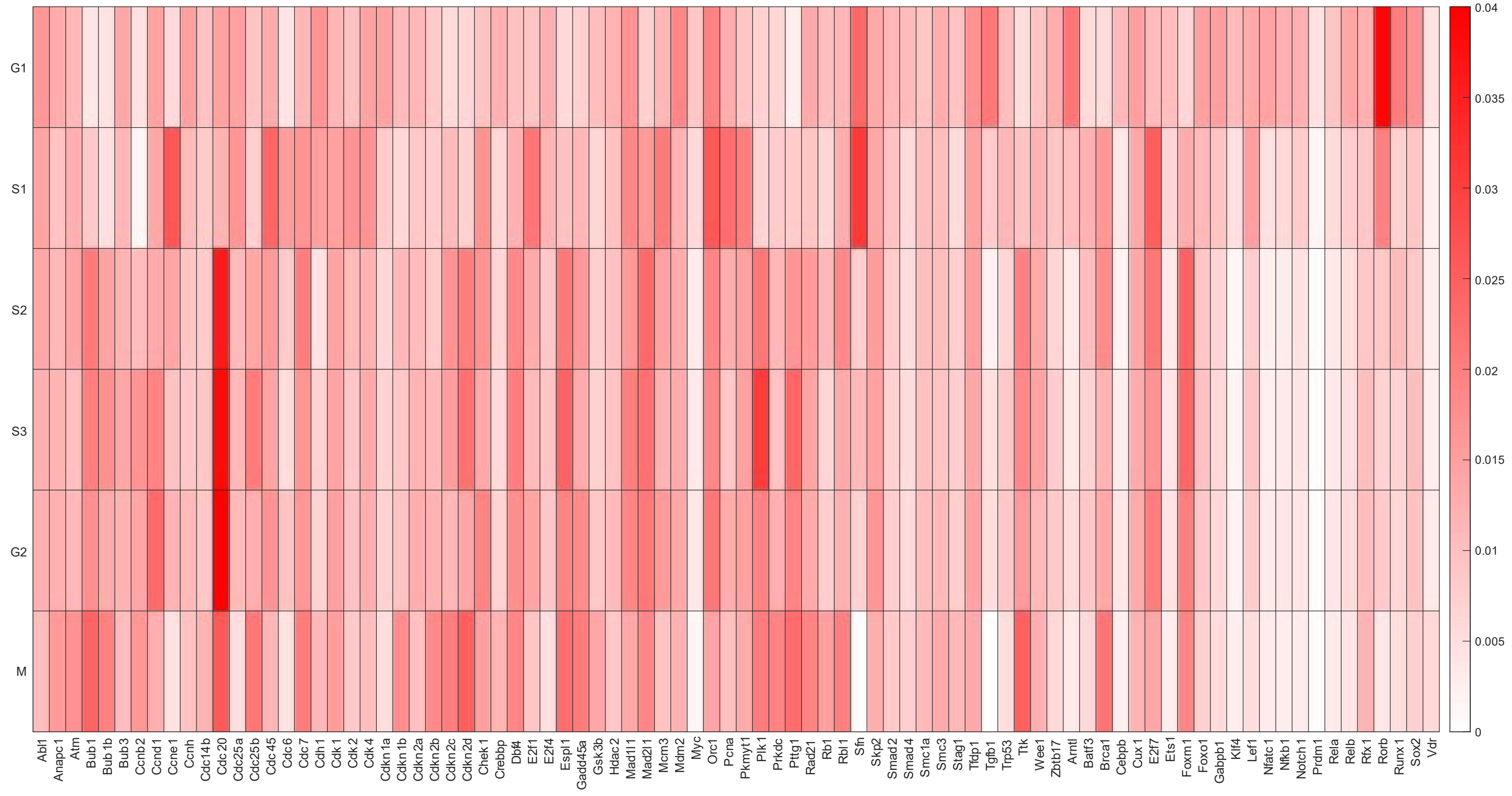


*Figure S5: Stage-specific cybernetic weights.*

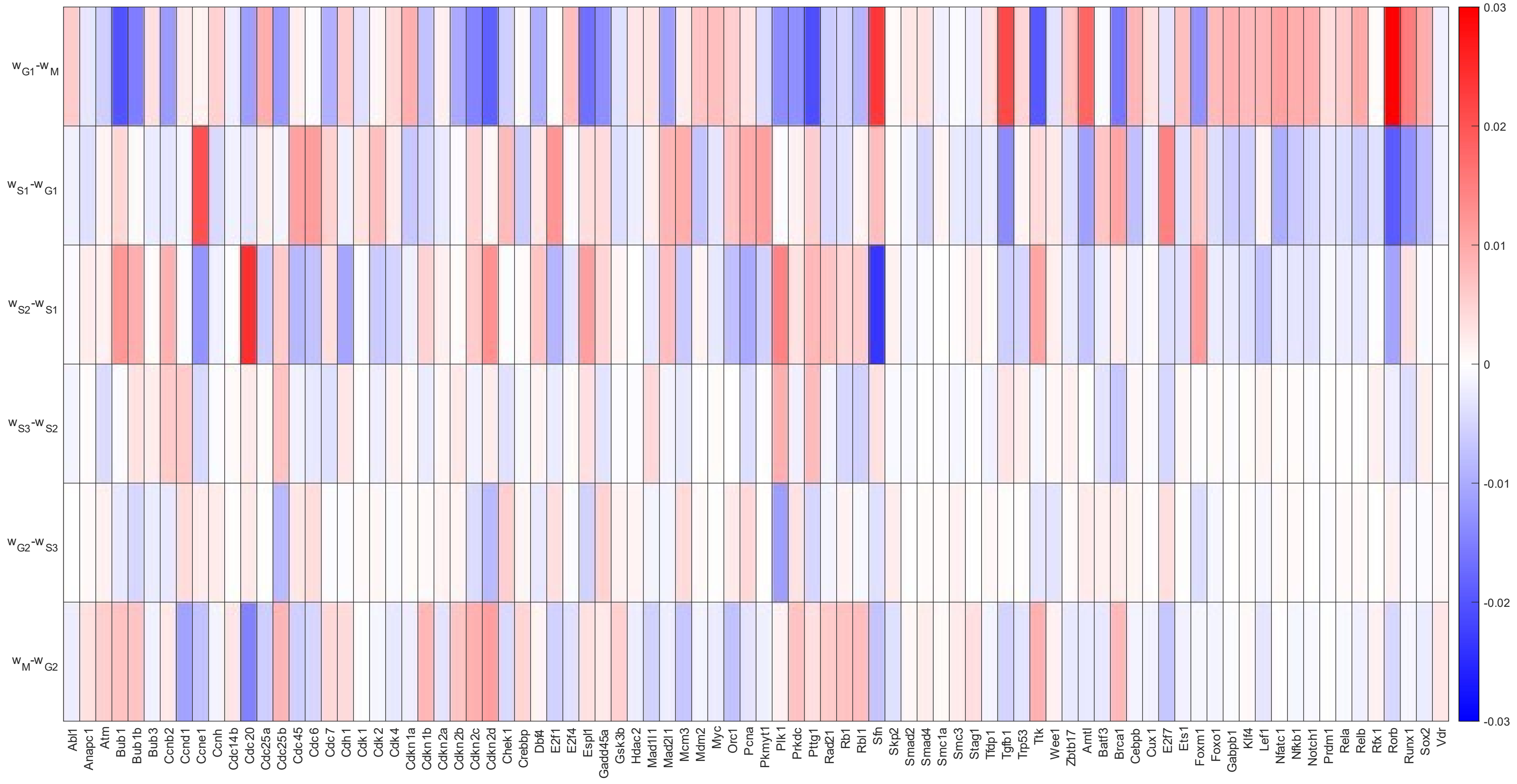


*Figure S6: Cybernetic weight differences between adjacent stages.*

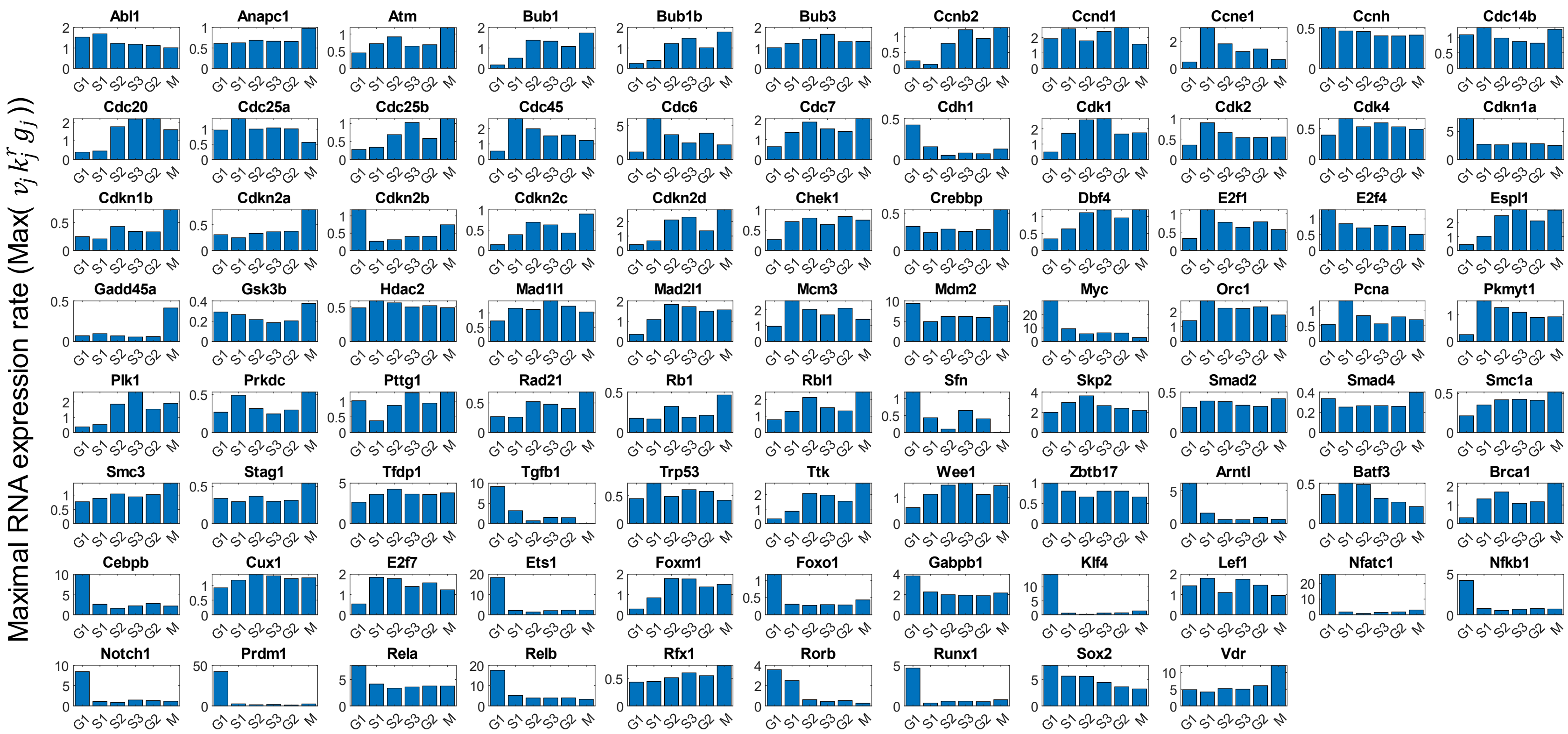


*Figure S7: Stage-specific Maximal RNA expression rate (Max($v_j k_j^r g_j$))*
